# Dynamic spreading of chromatin-mediated gene silencing and reactivation between neighboring genes in single cells

**DOI:** 10.1101/2021.11.04.467237

**Authors:** Sarah Lensch, Michael H. Herschl, Connor H. Ludwig, Joydeb Sinha, Michaela M. Hinks, Adi Mukund, Taihei Fujimori, Lacramioara Bintu

## Abstract

In mammalian cells genes that are in close proximity are coupled transcriptionally: silencing or activating one gene can affect its neighbors. Understanding these dynamics is important for natural processes, such as heterochromatin spreading during development and aging, and when designing synthetic gene regulation. Here, we systematically dissect this process in single cells by recruiting and releasing repressive chromatin regulators at dual-gene synthetic reporters, and measuring how fast gene silencing and reactivation spread as a function of intergenic distance and configuration of insulator elements. We find that silencing by KRAB, associated with histone methylation, spreads between two genes within hours, with a time delay that increases with distance. This fast KRAB-mediated spreading is not blocked by the classical cHS4 insulators. Silencing by histone deacetylase HDAC4 of the upstream gene can also lead to downstream gene silencing, but with a days-long delay that does not change with distance. This slower silencing can sometimes be stopped by insulators. Gene reactivation of neighboring genes is also coupled, with strong promoters and insulators determining the order of reactivation. We propose a new model of multi-gene regulation, where both gene silencing and gene reactivation can act at a distance, allowing for coordinated dynamics via chromatin regulator recruitment.

## Introduction

Eukaryotic gene expression is regulated by chromatin regulators (CRs) and the histone and DNA modifications they read, write, and remove^1, 2^. Chromatin-mediated gene regulation is crucial in development, aging, and disease^3–5^; it is also important in synthetic biology, where precise control of gene expression is necessary^6, 7^. Site-specific recruitment of CRs is a common method of regulating gene expression in synthetic systems. CRs modulate gene expression with varying kinetics and can establish long-term epigenetic memory through positive feedback mechanisms^8–11^. However, positive feedback also enables spreading of epigenetic effects beyond the target locus, which can lead to undesirable changes in global gene expression. Neither the temporal dynamics nor the spatial extent of this process are well characterized. Here, we seek to understand the effects of intergenic distance and insulators on the dynamics of spreading of gene silencing and activation between two neighboring genes. This understanding will be useful for building more robust synthetic systems and epigenetic therapies.

To understand the effects of intergenic distance and insulators on spreading, we studied the dynamics of spreading of gene silencing after recruitment of different types of CRs to a dual-gene reporter (Figure 1A-B). This reporter consists of two fluorescent genes separated by increasing DNA distances or insulator elements derived from the chicken hypersensitivity site 4 (cHS4)^12–14^. At this reporter, we recruited two types of chromatin regulators: Kruppel associated box (KRAB) and Histone deacetylase 4 (HDAC4).

**Figure 1.**
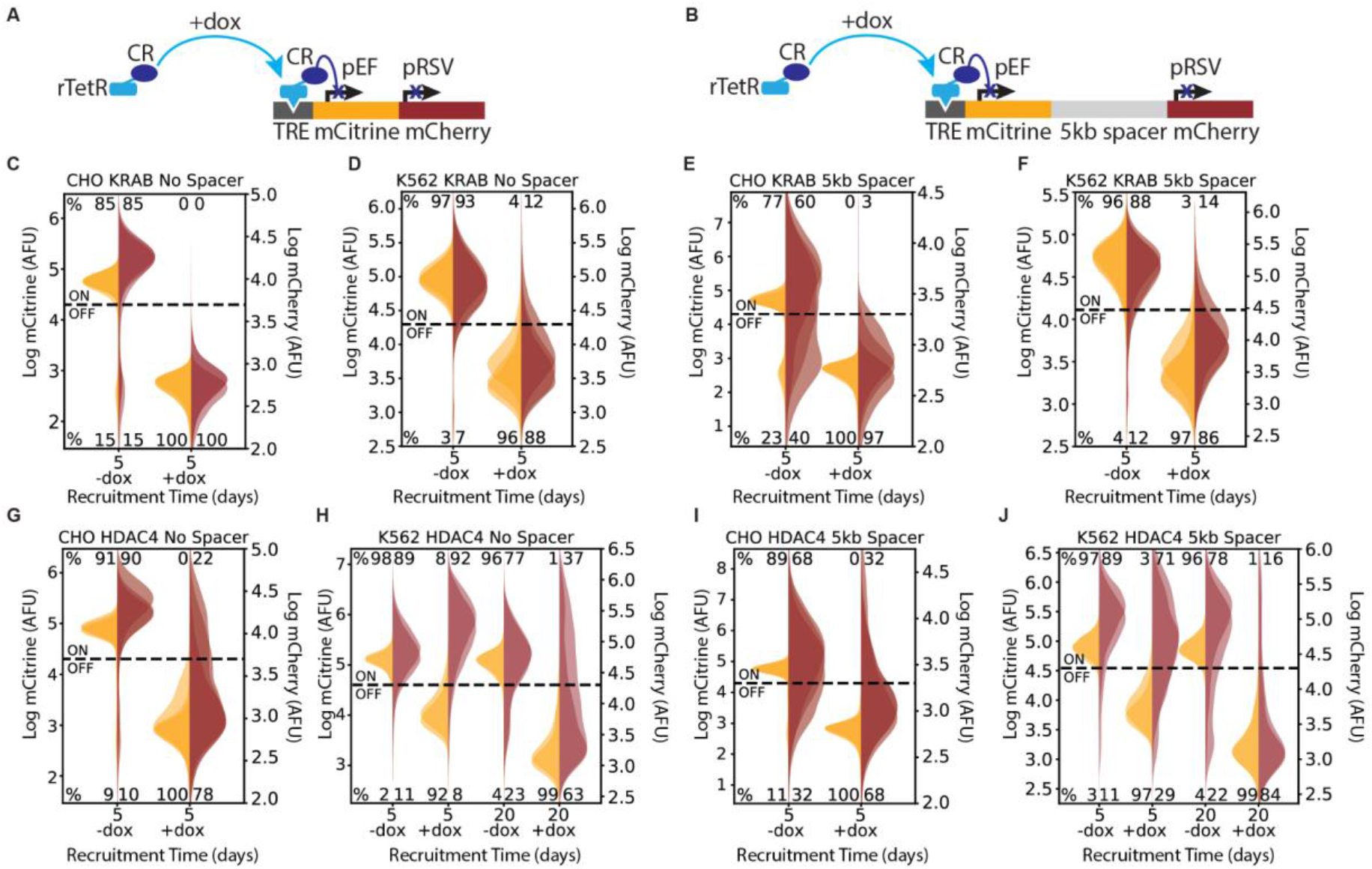
Recruitment of chromatin regulators to synthetic dual-fluorescent reporters results in transcriptional silencing of both genes. **(A, B)** Recruitment of a chromatin regulator (dark blue oval) via addition of dox allows for the binding of the rTetR-CR fusion to TRE (Tet Responsive Element, dark grey), upstream of a dual-gene reporter expressing mCitrine (yellow) and mCherry (red) separated by either **(A)** no spacer or **(B)** 5kb of lambda DNA (gray). **(C-J)** Fluorescence distributions of mCitrine and mCherry measured by flow cytometry either without CR recruitment (-dox) or after 5 or 20 days of recruitment (+dox) of either KRAB or HDAC4 at the NS or 5kb reporters in either CHO-K1 or K562 as indicated in each title. Percentages of cells ON (high-fluorescence, top) or OFF (low-fluorescence, bottom) are calculated based on a threshold (dotted line). Data from independent clonal cell lines for CHO-K1 or biological replicates of multiclonal populations for K562 are shown as overlaid semi-transparent distributions (n=3). **Figure 1 - figure supplement 1.** Reporter constructs used in different cell lines for analyzing spreading of transcriptional changes. **Figure 1 - figure supplement 2.** Transcriptional run-on from pEF over pRSV.

The Kruppel associated box (KRAB) from zinc finger 10 is a commonly used repressive domain that is associated with spreading of heterochromatin and epigenetic memory through positive feedback mechanisms^8, 15–22^. KRAB-mediated gene silencing operates through recruitment of cofactors, including KAP1, HP1, and SETDB1, that read and write the repressive histone modification, histone 3 lysine 9 trimethylation (H3K9me3), creating a positive feedback loop that establishes stable gene silencing^9^. This type of feedback also enables spreading of epigenetic effects beyond the target locus^19, 20, 23^.

Histone deacetylase 4 (HDAC4) is a repressive CR that functions primarily through removal of histone acetylation^24, 25^. Unlike KRAB, however, HDAC4 does not exhibit positive read-write feedback. In addition, our previous studies show that removal of HDAC4 after silencing leads to quick and complete reactivation of the gene^8^. Due to lack of positive feedback and memory, we hypothesized that spreading of silencing beyond the target locus would not occur upon HDAC4 recruitment.

We used single-cell time-lapse microscopy and flow cytometry to monitor gene expression during and after KRAB or HDAC4 recruitment to the upstream gene. We found that transcriptional silencing of the upstream gene by recruitment of either KRAB or HDAC4 affects the downstream gene even when separated by up to 5kb of distance or with different insulator arrangements. For KRAB, the time required for spreading to occur is short and largely distance-dependent. For HDAC4 however, the delay time between silencing of the two genes is longer, does not increase with intergenic distance, and can be influenced by insulators. We used these findings to form a kinetic model that leverages information from changes in histone modifications to understand the dynamics of gene expression during both silencing and reactivation. Our results show that targeted transcriptional silencing affects neighboring genes and provide insight for designing novel synthetic systems in mammalian cells.

## Results

### Spreading of transcriptional silencing between genes

To study the spreading of transcriptional silencing and chromatin modifications, we engineered a set of mammalian cell lines containing two neighboring fluorescent reporter genes and a CR, with each gene driven by constitutive promoters (Figure 1A-B, Figure 1–figure supplement 1). The addition or removal of doxycycline (dox) induces the recruitment or release, respectively, of the rTetR-fused CRs upstream of the first reporter gene, enabling precise control over the duration of recruitment (Figure 1A-B). To finely probe the effect of intergenic distance on spreading dynamics, we separated the two fluorescent genes by various spacer lengths. Specifically, we separated the genes with no spacer (NS) (Figure 1A) or 5kb lambda phage DNA (5kb) (Figure 1B), which was used as a neutral spacer^26^. To mitigate variability due to genomic position, we site-specifically integrated our reporters in two cell types: in CHO-K1 (Chinese Hamster Ovarian) at the phiC31 integration site on the multi-integrase human artificial chromosome (MI-HAC)^27^ and in human K562 at the AAVS1 safe harbor locus on chromosome 19 (Figure 1–figure supplement 1)^28^.

To investigate the spreading of transcriptional silencing, we first recruited KRAB upstream of the mCitrine gene for 5 days in CHO-K1 and K562 cell lines with the addition of dox (Figure 1A-B). Using flow cytometry to measure fluorescence intensity, we observed silencing of both the upstream mCitrine gene and the downstream mCherry gene in the NS (Figure 1C-D) and 5kb (Figure 1E-F) reporter lines for both cell types. This observation is consistent with previous reports of spreading of silencing upon KRAB recruitment^20, 23^. Surprisingly, HDAC4 recruitment also resulted in silencing of both genes (Figure 1G-J), albeit with reduced strength: only 68-78% of CHO-K1 cells silenced mCherry with HDAC4 (Figure 1G&I) compared to 97-100% with KRAB (Figure 1C&E). Moreover, in K562 cells, HDAC4-mediated silencing of mCherry was much slower, taking up to 20 days, compared to 5 days in CHO-K1 cells (Figure 1H&J). These results show for the first time that silencing mediated by a histone deacetylase can also extend to nearby genes.

Additionally, we noticed that in CHO-K1 cells in the absence of dox, mCherry expression is lower and has a wider distribution in the 5kb reporter compared to NS (Figure 1E&I vs. C&G). This observation suggests that the two promoters interfere, as observed before^29^, in a distance-dependent manner. Surprisingly, in K562 we saw an increase in pRSV-mCherry expression after 5 days of dox-mediated HDAC4 recruitment and pEF-mCitrine silencing (Figure 1H). We found that this transcriptional interference between pEF and pRSV results from transcriptional run-on from the pEF promoter which is resolved upon mCitrine silencing (Figure 1H, Figure 1–figure supplement 2A-B, Supplemental Text). Despite this type of transcriptional interference, we can still measure the silencing dynamics of the downstream gene.

### Changes in histone modifications upon transcriptional silencing of neighboring genes

To see if the changes in gene expression were accompanied by changes in chromatin modifications, we used CUT&RUN^30, 31^ to map activating and repressive histone modifications. Recruitment of KRAB for 5 days in K562 cells at the 5kb reporter resulted in loss of the activating histone 3 lysine acetylation (H3Kac) and a strong increase in the repressive modification H3K9me3 across the recruitment site, both fluorescent protein genes, and the 5kb spacer (Figure 2A). This increase in H3K9me3 and decrease in H3Kac was also observed in the NS reporter line (Figure 2–figure supplement 1A) and was accompanied by a concomitant depletion of histone 3 lysine 4 trimethylation (H3K4me3), which is associated with active genes, in both reporter lines (Figure 2–figure supplement 1A-B). Moreover, we observed an increase in H3K9me3 beyond the AAVS1 integration site into neighboring genes (Figure 2B), corresponding to a region of enrichment of approximately 50-60 kb beyond the reporter (Figure 2–figure supplement 1C-D). This too was accompanied by a depletion of H3Kac and H3K4me3 in the immediate vicinity of the AAVS1 integration site. The magnitude of histone modification changes in this vicinity was comparable between the NS and 5kb reporter lines (Figure 2–figure supplement 1E). We observed similar changes in chromatin modifications in CHO-K1 cells after 5 days of KRAB recruitment to the 5kb reporter (Figure 2–figure supplement 2A), showing that these changes in histone modifications coincide with the spreading of transcriptional silencing at the reporter in different cell lines.

**Figure 2.**
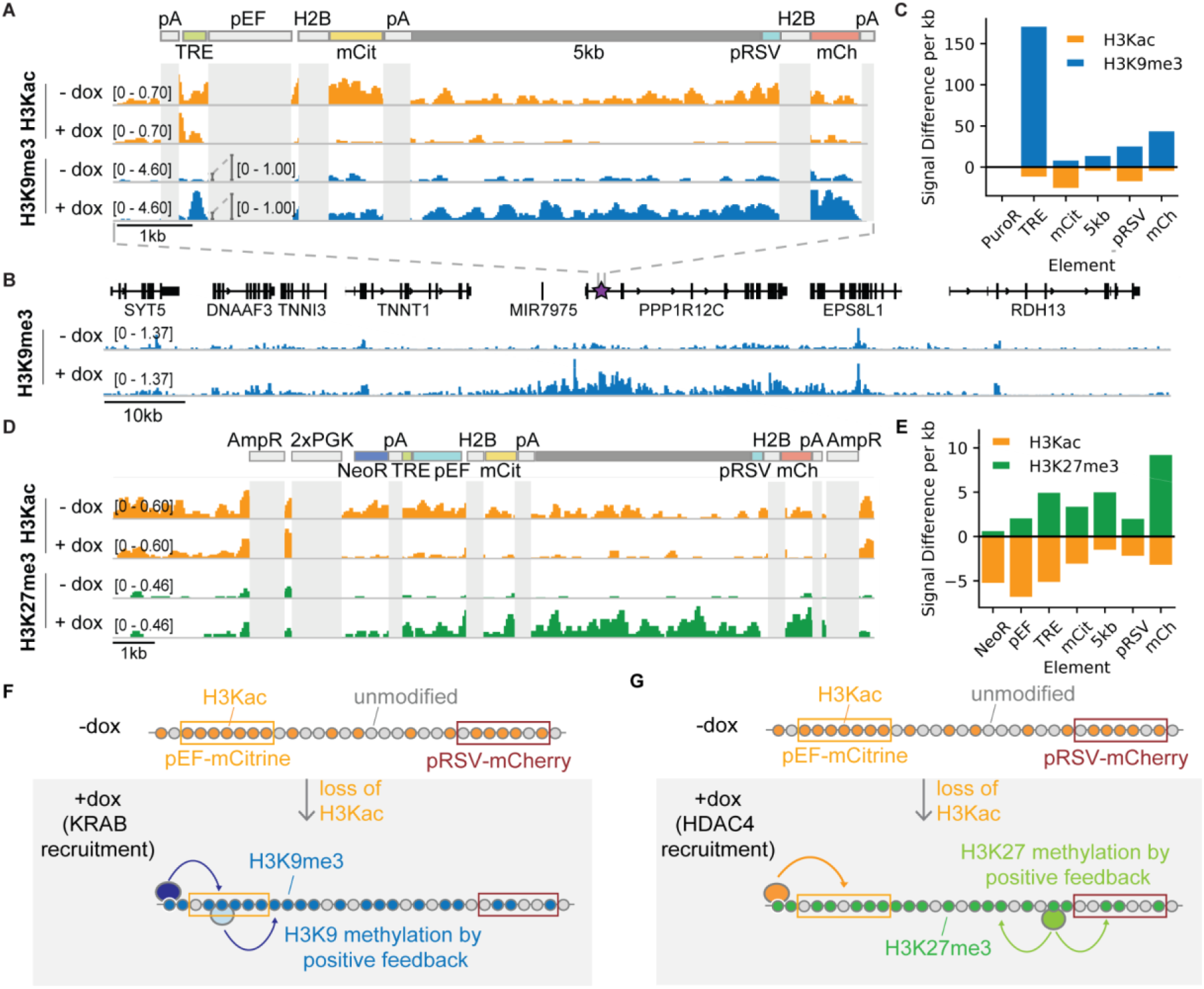
CUT&RUN data shows changes in histone modifications at silenced genes. **(A)** Genome browser tracks showing counts per million (CPM)-normalized reads after CUT&RUN for histone 3 acetyl-lysine (H3Kac) and histone 3 lysine 9 trimethylation (H3K9me3) with (+dox) and without (-dox) recruitment of rTetR-KRAB to the 5kb-spacer reporter in K562 cells. Non-unique regions resulting in ambiguous alignment, including pEF, H2B, and polyA tails (pA), are masked in light gray. **(B)** Genome browser tracks of H3K9me3 with (+dox) and without (-dox) recruitment of rTetR-KRAB, looking at the surrounding locus where the 5kb reporter is integrated in cells (purple star within first intron of PPP1R12C). This snapshot does not include an in-situ representation of the reporter, which instead was appended to the end of the reference genome to preserve gene annotations. **(C)** Quantification of signal difference for H3K9me3 and (H3Kac) with (+dox) and without (-dox) recruitment of KRAB for 5 days to the 5kb reporter in K562 cells. **(D)** Genome browser tracks showing CPM-normalized reads after CUT&RUN for H3Kac and histone 3 lysine 27 trimethylation (H3K27me3) with (+dox) and without (-dox) recruitment of rTetR-HDAC4 to the 5kb-spacer reporter in K562 cells. Non-unique regions resulting in ambiguous alignment, including AmpR, PGK, H2B, and polyA tails (pA), are masked in light gray. **(E)** Quantification of signal difference for H3Kac and H3K27me3 with (+dox) and without (-dox) recruitment of HDAC4 for 5 days to the 5kb reporter in K562 cells. **(F)** Distance-dependent delay of transcriptional silencing between two genes via recruitment of KRAB leads to loss of H3Kac and gain of H3K9me3 across the dual-gene reporter through both DNA looping from rTetR-KRAB as well as positive feedback loops for spread of methylation. **(G)** Distance-independent delay of transcriptional silencing between two genes via recruitment of HDAC4 leads to a loss of H3Kac as well as a gain of H3K27me3 across the reporter through positive feedback loops for spread of methylation. **Figure 2-figure supplement 1.** Changes in chromatin modifications at the two-gene reporter and surrounding AAVS1 locus after recruitment of KRAB for five days in K562 cells. **Figure 2 - figure supplement 2.** Recruitment of KRAB or HDAC4 for five days in CHO-K1 cells induces changes in chromatin modifications at the two-gene reporters.

Recruitment of HDAC4 for 5 days in CHO-K1 cells to the 5kb and NS reporters showed a decrease in acetylation levels, and no change in H3K9me3 (Figure 2D-E, Figure 2–figure supplement 2B-C), as expected. However, surprisingly histone 3 lysine 27 trimethylation (H3K27me3) was also detected throughout the 5kb (Figure 2D-E) and the NS reporters (Figure 2– figure supplement 2B). This repressive modification, which is associated with polycomb repressive complex 2 (PRC2), has not been reported in association with HDAC4, and was not observed after HDAC4 recruitment at a pEF-mCitrine reporter flanked by insulators in the same locus of CHO-K1 cells ^8^. Like H3K9me3 (Figure 2E), H3K27me3 has reader-writer positive feedback ^10^ that can lead to spreading of these chromatin modifications (Figure 2F). This observation suggests that silencing of the RSV promoter is indirectly due to PRC2 action, but it can only take place once HDAC4 removes acetylation at the locus. This system allows us to study the temporal and distance dependence dynamics of this type of indirect spreading of silencing, where silencing of one gene opens the door for other repressive complexes to silence neighboring genes (Figure 2E). We can compare these dynamics with those associated with direct silencing by KRAB, where repressive modifications that are associated with the recruited chromatin regulator spread across the locus (Figure 2F).

### Dynamics of spreading of transcriptional silencing— delay between two genes

We measured the spreading dynamics of transcriptional silencers at finer temporal resolution by first taking time-lapse movies of CHO-K1 cells with the NS and 5kb spreading reporters (Movies S1-S4). Cells were imaged every 20 mins for 4-5 days and cumulative fluorescence traces were used to determine transcriptional silencing times after dox addition (Figure 3A, Figure 3–figure supplement 1). Upon KRAB recruitment, we observed only a slight delay of 2.3±3.1 hours between mCitrine and mCherry silencing in the NS reporter and a longer delay of 12.3±9.8 hours in the 5kb reporter (Figure 3B&C, Figure 3–figure supplement 1A&B). In contrast, spreading of HDAC4-mediated transcriptional silencing was slower, and no appreciable difference in silencing delays was observed between reporter lines (NS: 37+/- 21 hours; 5kb: 32+/-19 hours) (Figure 3D&E, Figure 3–figure supplement 1C&D). Taken together, these data suggest that KRAB-mediated spreading of transcriptional silencing is distance-dependent in CHO-K1 cells while HDAC4 mediated spreading is distance-independent at these length scales.

**Figure 3.**
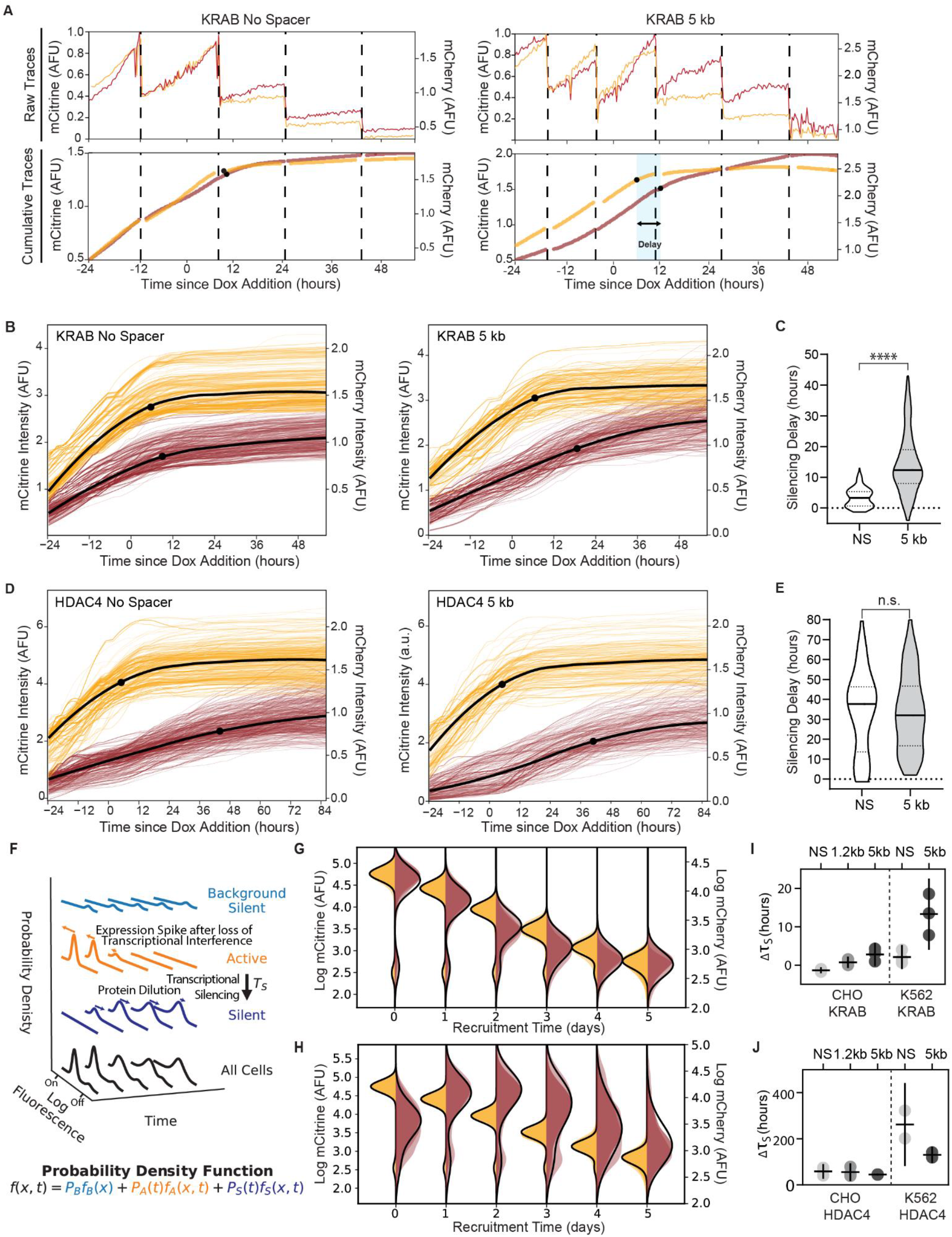
Single-cell data measures delay in transcriptional silencing of the two genes. **(A)** (Top) Example raw traces measured by time-lapse microscopy showing total fluorescence of mCitrine (yellow) and mCherry expression (red) as a function of time in an individual cell lineage in CHO-K1 with the NS reporter (left) and 5kb reporter (right). Dotted lines denote cell divisions. Recruitment of KRAB starts 24 hours by dox addition. (Bottom) Cumulative single-cell traces stitched across cell divisions showing estimated silencing time points (black dots, Materials and Methods) and silencing delay (blue shading). **(B)** Cumulative single**-**cell traces of mCitrine (yellow) and mCherry (red) in CHO-K1 cells with NS reporter (left, n=296) and 5kb reporter (right, n=218) with the mean trace (black curves) and median silencing times (dots) as a function of time since recruitment of KRAB by dox addition at time 0. **(C)** Distribution of delay times between silencing of mCitrine and mCherry in individual cells as shown in (A) after recruitment of KRAB in (NS reporter median silencing delay = 3.3 hours; 5kb reporter median silencing delay = 12.3 hours; statistically significant difference by Welch’s unequal variances T-test). **(D)** Cumulative single-cell traces as a function of time relative to HDAC4 recruitment (NS, n= 211; 5kb, n=291), as in (B). **(E)** Distribution of delay times after recruitment of HDAC4 (NS reporter median silencing delay = 38 hours; 5kb reporter median silencing delay = 32 hours; no statistically significant difference by Welch’s unequal variances T-test). **(F)** A probabilistic model consisting of three states: background silent (light blue), active (orange), silent (dark blue). Each state has its own weight and distribution, and all states are summed to a final probability density function (black) that describes the probability ***f*** of finding a cell with fluorescence ***x*** at time ***t***. This model is used to fit daily flow cytometry data to extract transcriptional silencing times (***τ***_***S***_) upon CR recruitment, while taking into account: stochastic transitions of cells from the active to the silent state upon CR-mediated silencing, spike in expression after loss of transcriptional interference, and mRNA and protein degradation and dilution (Supplementary Text). **(G, H)** Overlaid daily distributions of mCitrine (transparent yellow) and mCherry (transparent red) fluorescence from flow cytometry during recruitment of **(G)** KRAB and **(H)** HDAC4 with average model fit (black line) (n=3). **(I)** Silencing delays between mCitrine and mCherry after KRAB recruitment extracted from daily flow cytometry time-courses using the model in (F) for different spreading reporters: NS, 1.2kb, 5kb. Each dot represents a clone for CHO-K1 (left) and a biological replicate for K562 (right); horizontal bar is mean delay, vertical bar is 90% confidence interval from the fit estimated using the t-distribution (n=3). **(J)** Silencing delays between mCitrine and mCherry after HDAC4 recruitment (CHO-K1, n=3 clones; K562 n=2 biological replicates); same notation as (H). **Figure 3 - figure supplement 1.** Example images and single-cell analysis of silencing dynamics from time-lapse microscopy of CHO-K1 cells. **Figure 3 - figure supplement 2.** Dynamics of silencing measured by flow cytometry and fit by gene expression model. **Figure 3 - figure supplement 3.** Steady-state expression in the absence of dox in CHO-K1 versus K562 cells.

Since time-lapse microscopy measurements are limited to hundreds of cells and hard to extend to non-adherent cell lines like K562, we developed an alternative approach to extracting the delay times between silencing of the two reporter genes from flow cytometry data. We developed a mathematical model that describes the evolution of mCitrine and mCherry fluorescence distributions after CR recruitment (Figure 3F, Materials and Methods). We used our daily flow cytometry measurements of fluorescence distributions to fit silencing times for each gene (Figure 3G-J), along with other parameters associated with gene expression including a spike in mCherry mRNA production due to transcriptional interference (as shown by qPCR in Figure 1– figure supplement 2C&D) and mRNA and protein degradation and dilution (Materials and Methods). This approach allowed us to estimate the delay between mCitrine and mCherry silencing, Δ***τ***_s_, at the gene rather than protein level for all spreading reporters in CHO-K1 and K562 with different CRs recruited (Figure 3G-J, Figure 3–figure supplement 2, Table S2). Overall, the delay between mCitrine and mCherry silencing are similar between movie traces and fits of the cytometry data (Figure 3C&E vs. I&J), especially if we consider the additional time necessary for mRNA degradation of about ∼4 hours^8^ included in the flow fits (Materials and Methods).

These delays show similar trends with distance in CHO-K1 and K562 cells for a given CR. KRAB recruitment results in silencing delays that increase with intergenic distance (Figure 3I). HDAC4 recruitment results in delays that are not statistically different among different distances in either cell type, but show a trend where delay decreases with increased distance in K562 from around 10 days in the NS reporter to around 5 days in the 5kb reporter (Figure 3J). The observation that the delays of pRSV-mCherry silencing do not increase with distance (Figure 3E&J) suggests that its silencing initiates at the RSV promoter, most likely by the action of PRC2 (Figure 2D, E&G).

Silencing spreads slower in K562 versus CHO-K1 cells after both KRAB and HDAC4 recruitment (Figure 3I vs J), consistent with the 5 days results (Figure 1). This is most likely because the pRSV promoter is stronger in K562 cells, as indicated by higher mCherry expression compared to CHO-K1 cells for both the NS and 5kb reporters (Figure 3–figure supplement 3). These small differences in temporal dynamics are likely due to differences in context of the reporters, both different genomic locus and cell type.

### The role of the cHS4 insulators in spreading of transcriptional silencing between genes

Insulators are sequences shown to prevent transgene silencing and believed to stop spreading of repressive chromatin modifications^32^. The most commonly used insulator is cHS4; most of its insulator activity has been attributed to a 250bp core region^13, 33–36^ that is associated with increased histone acetylation^37, 38^. To measure the role of the cHS4 insulator on blocking the spreading of targeted gene silencing, we built four different insulator configurations: a “single insulator” between the two genes using either the full length 1.2kb cHS4 (SH) or its 250bp core region (SC) (Figure 4A), or insulators flanking the mCitrine gene, referred to as “double insulator” (Figure 4B), with either full cHS4 (DH) or core region (DC). This double configuration is commonly used in mammalian cell engineering to prevent background silencing of transgenes due to position effect variegation^12^.

**Figure 4.**
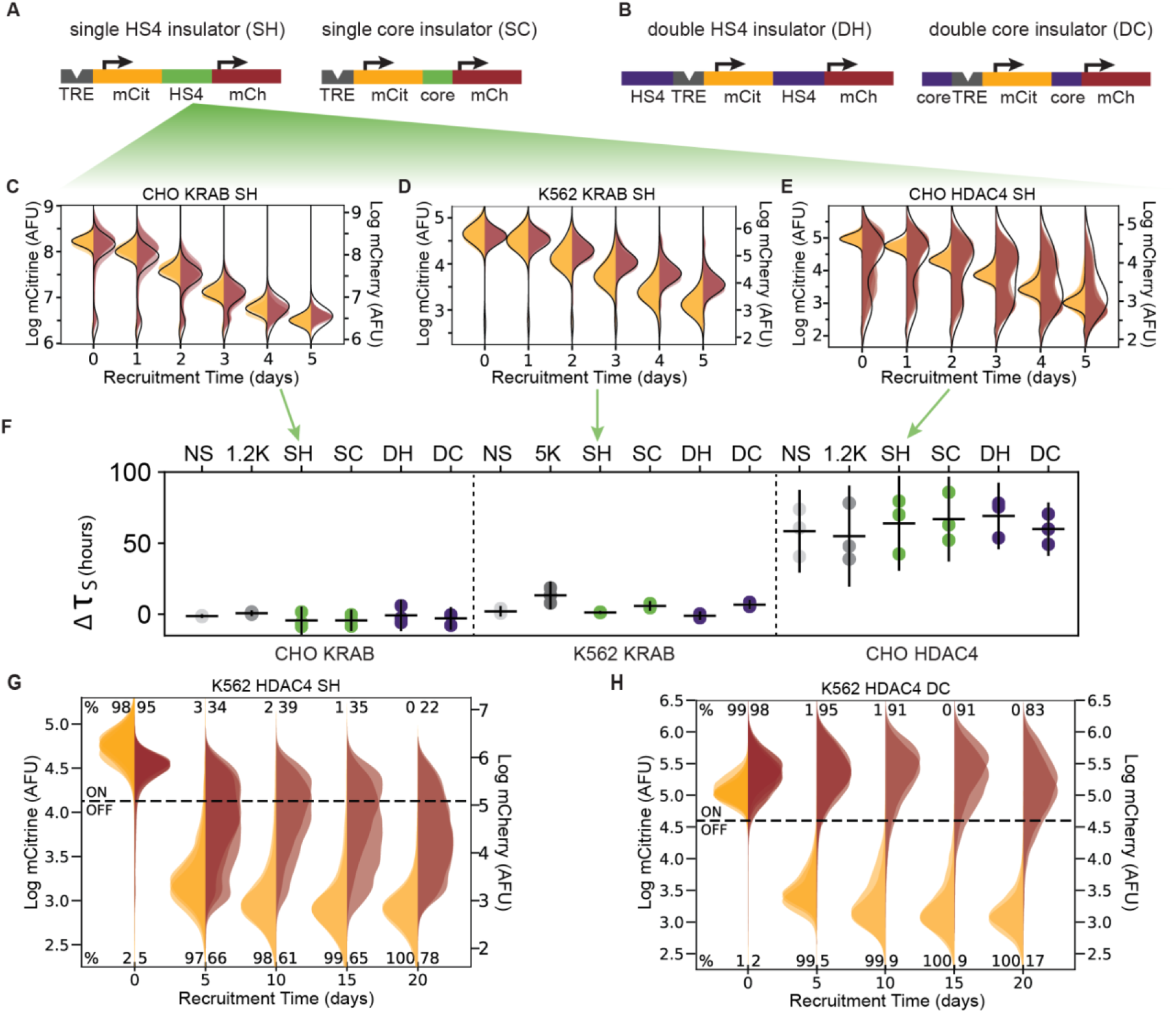
The role of the cHS4 insulators in spreading of transcriptional silencing across genes. **(A)** Single insulator geometries between mCitrine (mCit) and mCherry (mCh) fluorescent genes, with full 1.2kb HS4 insulator (SH, left) and 250 bp core insulator (SC, right). **(B)** Double insulator geometries with full length 1.2kb HS4 insulator (DH, left) and 250 bp core insulator (DC, right) flanking the TRE-pEF-mCitrine region. **(C-E)** Overlaid replicates of daily distributions of mCitrine (yellow) and mCherry (red) fluorescence from flow cytometry during recruitment of **(C)** KRAB in CHO-K1, (D) KRAB in K562, or **(E)** HDAC4 in CHO-K1, to SH reporters with average model fit (black line) (n=3). **(F)** Silencing delay times between mCitrine and mCherry in different insulator reporters after chromatin regulator recruitment for 5 days (each dot is a replicate, horizontal bar is mean delay, vertical bar is 90% confidence interval estimated using the t-distribution). **(G-H)** Overlaid replicates of daily distributions of mCitrine (yellow) and mCherry (red) fluorescence from flow cytometry during extended recruitment of HDAC4 to **(G)** SH reporter or **(H)** DC reporter in K562 (n=3). **Figure 4 - figure supplement 1.** The effect of all insulator configurations on spreading dynamics of transcriptional silencing in CHO-K1 and K562. **Figure 4 - figure supplement 2.** Insulators attenuate spreading of silencing via HDAC4 in K562. **Figure 4 - figure supplement 3.** Dynamics of spreading upon weaker gene targeting insulator reporters with HDAC4 at lower dox concentrations in CHO-K1. **Figure 4 - figure supplement 4.** Insulators do not block spreading of silencing with weaker gene targeting at lower dox concentrations. **Figure 4 - figure supplement 5.** Insulators can prevent background silencing of reporter genes.

Surprisingly, no insulator configuration was capable of stopping KRAB-mediated spreading of silencing in either CHO-K1 or K562 cells (Figure 4C-D, Figure 4–figure supplement 1A-B). Moreover, insulators do not delay KRAB-mediated silencing either: the estimated delay times in the insulator reporters are close to the delays in reporters with spacers (Fig 3F, Table S2). Therefore, these insulator configurations do not have a strong effect on silencing mediated by KRAB recruitment.

In CHO-K1 cells, no insulator configuration was able to inhibit spreading of silencing upon HDAC4 recruitment (Figure 4E, Figure 4–figure supplement 1C), and the small fraction of cells where mCherry is not silenced by day 5 of recruitment is similar to the spacer reporters (Figure 3– figure supplement 2B). Also similar to the spacer reporters, the fitted delay times ranged from 42-85 hours (Figure 4F right, Figure 4–figure supplement 1C). These delay times fall within the 90% confidence interval of the 1.2 kb lambda spacer (Table S2). Therefore, insulators do not appear to have a strong effect on HDAC4-mediated silencing at our reporters in CHO-K1 cells.

However, insulators do appear to attenuate HDAC4-mediated mCherry silencing in K562 cells, even with 20 days of recruitment (Figure 3G&H, Figure 4–figure supplement 2). Rather than seeing complete silencing of the mCherry reporter, we see a broadening of the mCherry fluorescence distribution with the majority of cells expressing lower levels of mCherry than the population without dox, both at 5 and 20 days of HDAC4 recruitment to insulator constructs in K562 (Figure 4G, Figure 4–figure supplement 2). An exception to this observation is the DC configuration, where the mCherry distribution remains in the ON range even after 20 days of HDAC4 recruitment (Figure 4H). The lack of complete mCherry silencing and broader mCherry distribution is in contrast to the silencing seen at the NS and 5kb reporters after 20 days of HDAC4 recruitment in K562 (Figure 1H&J). These results suggest that the insulators can help the downstream gene remain active in conditions where its silencing is already slow, such as after indirect silencing by HDAC4 recruitment in K562.

To test if insulators can block weaker gene targeting by HDAC4 in CHO-K1 cells, we performed silencing experiments at non-saturating dox concentrations. At lower levels of dox, silencing of the two genes is slower; fits of daily flow cytometry data show silencing delay times that decrease upon increasing dox concentrations for all insulator configurations (Figure 4–figure supplement 3). At lower dox concentrations, fewer cells silence mCitrine by the end of 5 days, but those that do are likely to also silence mCherry for both HDAC4 and KRAB (Figure 4–figure supplement 4), showing that the insulators do not block spreading of silencing in CHO-K1 even with weaker CR targeting.

Although the cHS4 insulators do not generally prevent spreading of silencing during recruitment of CRs, the insulators are able to prevent spontaneous background silencing of the reporters in the absence of dox in CHO-K1 cells (Figure 4–figure supplement 5) consistent with previous transgene silencing reports^34, 39^. Overall, levels of mCherry expression are higher in the insulator constructs (Figure 3–figure supplement 3), and there is no transcriptional run-on mRNA across the SH insulator (Supplementary Text, Figure 1–figure supplement 2E-F), suggesting it helps terminate transcription and prevent transcriptional interference as shown before^40^.

### Reactivation of genes

To understand the dynamics of neighboring gene coupling during transcriptional activation, we investigated the order of gene reactivation and the degree of epigenetic memory in our various two-gene constructs. We first silenced our various reporters with different spacers or insulators for five days, then removed dox to release the CR and monitored gene expression every few days by flow cytometry (Figure 5A). In CHO-K1 cells, while both genes reactivated simultaneously in the NS reporter (Fig 5B, Figure 5–figure supplement 1A&B), mCitrine reactivated first in the 5kb reporter (Figure 5C, Figure 5–figure supplement 1C&D), both after release of KRAB or HDAC4. This pattern of a distance-dependent delay in reactivation between mCitrine and mCherry also holds for the insulator configurations for both KRAB and HDAC4: the full 1.2 kb long cHS4 configurations featured delayed mCherry reactivation (Figure 5D, Figure 5– figure supplement 2), compared to the more synchronous reactivation patterns observed in the 250 bp long core insulator configurations (Figure 5–figure supplement 2). The order of reactivation suggests that in CHO-K1 cells reactivation initiates at the stronger pEF promoter that drives mCitrine, and spreads in a distance-dependent manner to the weaker pRSV-mCherry gene.

**Figure 5.**
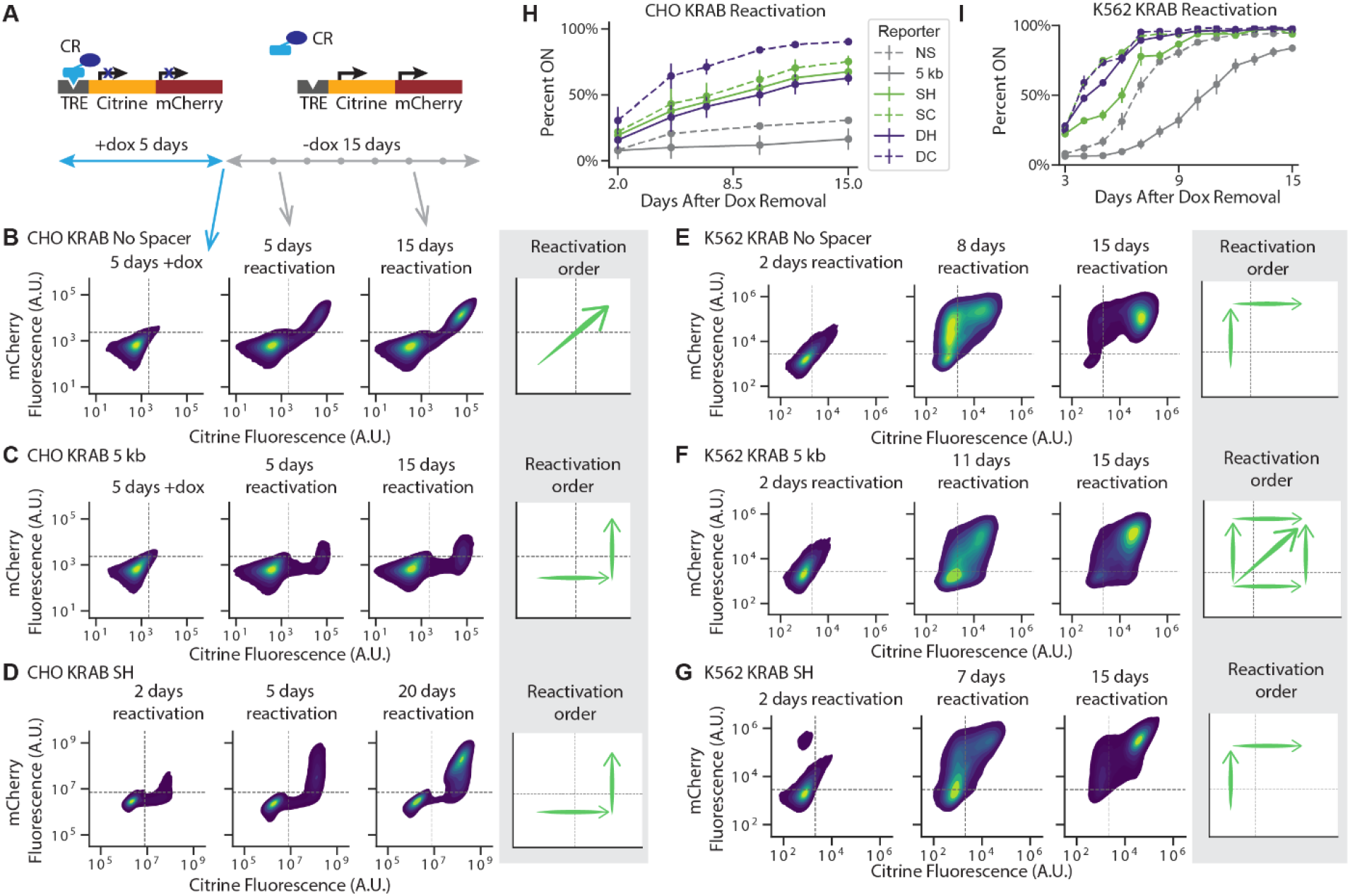
Reactivation of gene expression spreads between the two genes. **(A)** Schematic of experimental setup: each CR is recruited upstream of the mCitrine gene for 5 days. Recruitment is then stopped by removing dox and reactivation is monitored every few days by flow cytometry. **(B-G)** 2D density plots of mCitrine and mCherry fluorescence from flow cytometry show pattern of gene reactivation in : **(B)** CHO-K1 KRAB NS (n=3 clones), **(C)** CHO-K1 KRAB 5kb (n= 4 clones), **(D)** CHO-K1 KRAB SH (n=3), **(E)** K562 KRAB NS (n=3), **(F)** K562 KRAB 5kb (n=3), and **(G)** K562 KRAB SH(n=3). **(H-I)** Percent of cells in which at least one reporter gene reactivated over time after dox removal after KRAB release in **(H)** CHO-K1 and **(I)** K562. Replicates are from either independent clonal cell lines, where indicated, or from biological replicates of multiclonal populations. **Figure 5 - figure supplement 1.** Reactivation of gene expression in CHO-K1 NS and 5kb reporter lines. **Figure 5 - figure supplement 2.** Reactivation of gene expression in CHO-K1 insulator reporter lines. **Figure 5 - figure supplement 3.** Reactivation of gene expression in K562 cell reporter lines.

However, when looking at order of reactivation in K562 cells after release of KRAB, we found that for the NS reporter, mCherry reactivates first (Figure 5E), while with the 5kb reporter we observe three different scenarios in individual cells: either mCherry reactivates first, mCitrine reactivates first, or they reactivate together (Figure 5F). In the insulator constructs, mCherry reactivates before mCitrine in K562 cells (Figure 5G, Figure 5–figure supplement 3C). This change in reactivation pattern in K562 cells compared to CHO-K1 is likely due to the pRSV promoter being stronger in K562 (Figure 3–figure supplement 3).

Despite not preventing the spreading of silencing by KRAB, the insulators do play a role in the reactivation rate and level of memory. After release of rTetR-KRAB, reporters with insulators reactivated more rapidly compared to the NS and 5kb reporters, with the double core insulator exhibiting the highest degree of gene reactivation in both CHO-K1and K562 cells (Figure 5H). This is in line with the double core insulating elements also having the strongest insulating effect against mCherry silencing after HDAC4 recruitment in K562. Similarly, after removal of HDAC4 in CHO-K1 cells, insulators led both genes to reactivate to a higher extent (Fig 5I). These results show that the extent of epigenetic memory depends strongly not only on the chromatin regulator recruited, but also on the configurations of promoters and insulators at the target locus, with stronger promoters closely surrounded by core insulators diminishing memory.

### Model connecting chromatin states to gene expression dynamics

We used our gene dynamics and chromatin modifications data to develop a generalizable kinetic model that captures distance-dependent silencing associated with a tethered chromatin regulator and distills the roles of promoters and insulators as elements associated with high reactivation rates. In this model, each gene can be either active or silent, leading to four possible states in our two-gene reporters (Figure 6A). Each gene can transition between active and silent states with rates that depend on the distance between itself and other DNA elements that recruit chromatin regulators: the CR recruitment sites, promoters, and insulators. Under different experimental conditions, different rates in this kinetic pathway dominate, governing the transitions from one state to another. When KRAB is recruited upstream of pEF-mCitrine, silencing of the downstream pRSV-mCherry gene happens quickly (over hours) at a rate that decreases as the distance between pRSV-mCherry and pEF-mCitrine increases (Figure 6B, *k_s_KRAB_(d)*), suggesting the distance-dependent silencing rates induced by KRAB dominate over the activation rates associated with the promoters and insulators (Figure 6B, *k_a_* and *k_ins_* respectively).

**Figure 6.**
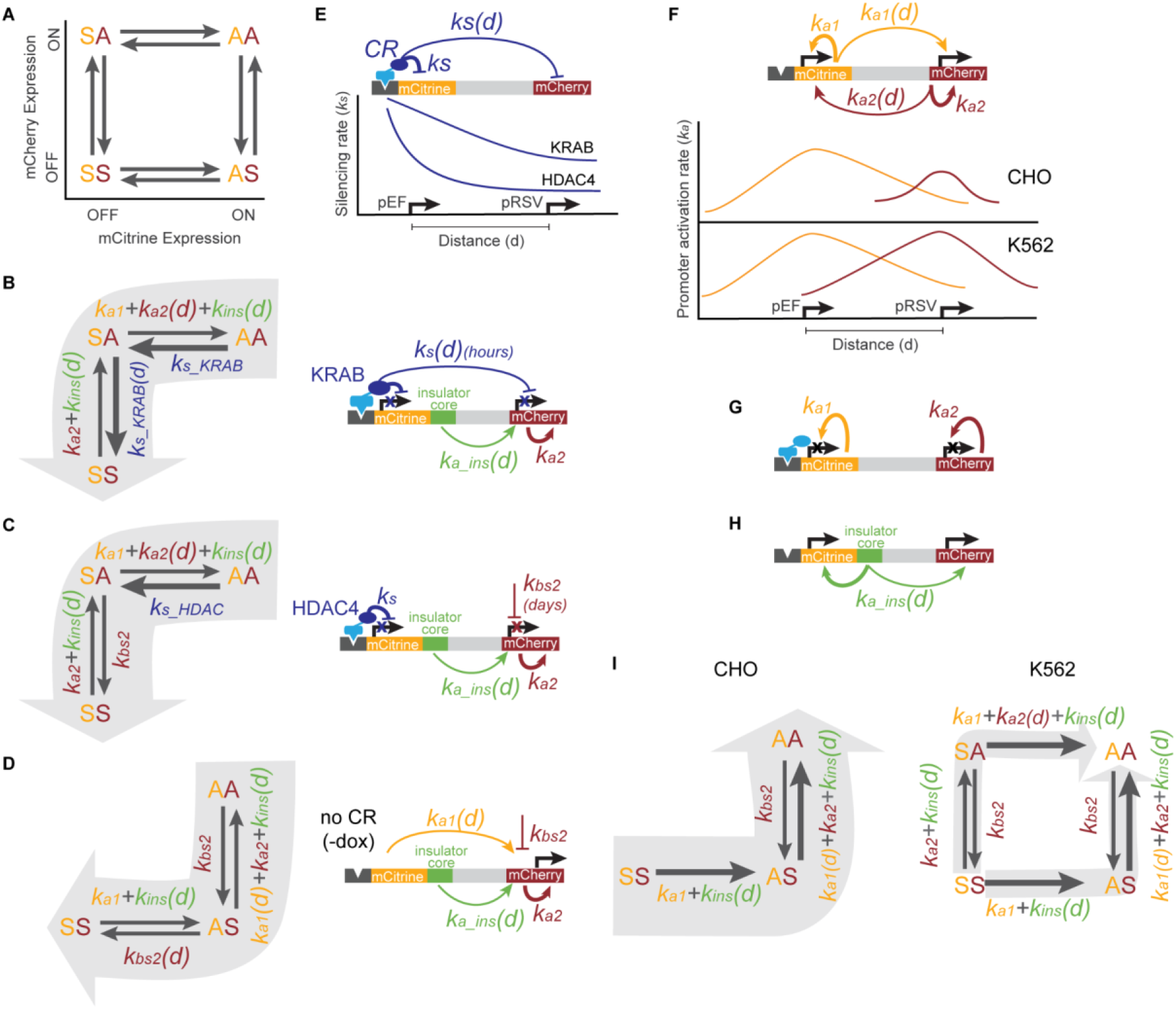
Model of multi-gene control coordinated by the action of CRs, promoters, and insulators. **(A)** Four states for a two-gene reporter, where the first letter (yellow) represents the mCitrine state and the second letter (red) represents mCherry state, as either active (A) or silent (S). Arrows represent the rates at which each gene is turned on or off. **(B)** During KRAB recruitment, cells transition from both genes active (AA) to both gene silent (SS), with citrine silencing first (SA intermediate state). The silencing rates of KRAB at the nearby pEF-mCitrine (*k_s_KRAB_*) and pRSV-mCherry (*k_s_KRAB_(d),* where *d* is the distance between the pEF and pRSV promoters) dominate over the activation rates of promoters (*k_a1_, k_a2_*) and insulators (*k_ins_(d),* where here *d* is the distance between the core insulator and a nearby promoter). **(C)** During HDAC4 recruitment, silencing of mCitrine (AA to SA) is driven by the silencing rate of HDAC4 (*k_s_HDAC4_*), while silencing of mCherry (SA to SS) is driven by background silencing rate (*k_bs2_*). Both the pRSV promoter (*k_a2_*) and insulator reactivation rates (*k_ins_)* can compete with pRSV-mCherry silencing. **(D)** Before CR recruitment (no dox), pEF as well as insulators can act from a distance on pRSV (*k_a1_(d), k_a_ins_(d)*) preventing background silencing of mCherry (*k_bs2_*). **(E)** KRAB can act on genes over a long distance (*k_s_(d)*), silencing both mCitrine and mCherry, while the range of HDAC4 silencing is much smaller (*k_s_*), only directly affecting mCitrine. **(F)** In the absence of CR recruitment, promoters can activate themselves (*k_a1_*, *k_a2_*), and maintain activity of genes at a distance (*k_a1_(d), k_a2_ (d)*). In CHO-K1, pRSV is weaker than in K562. **(G)** When genes are silenced, their promoters cannot act on neighboring genes, instead only reactivate themselves (*k_a1_*, *k_a2_*). **(H)** Insulators can maintain activity of the nearby genes and drive reactivation after CR-mediated silencing (*k_ins_(d)*). **(I)** Reactivation after silencing is driven by gene activation rates (*k_a1_*, *k_a2_*) along with insulators (*k_ins_(d)*) if present. In CHO-K1 cells where pEF is stronger than pRSV, mCitrine reactivates first followed by mCherry (left). However, in K562 where both promoters are equally strong, either gene can reactivate first (right).

However, when HDAC4 is recruited upstream of pEF-mCitrine, we observe delayed silencing of pRSV-mCherry (over many days, compared to hours in KRAB) at a rate that does not change as a function of the distance between the two genes. We hypothesize that pRSV-mCherry silencing is not due to the direct action of the HDACs recruited upstream of the pEF, but rather due to background silencing (Fig 6C, *k_bs2_*) of pRSV by endogenous polycomb complexes as indicated by the appearance of H3K27me3 (Figure 2D&E, Figure 2–figure supplement 2B&C).

Because we see much less pRSV-mCherry background silencing when the upstream pEF-citrine is active (in the absence of dox), we conclude that when pEF is active, it can increase activity of the downstream pRSV and prevent it from background silencing (Figure 6D, *k_a1_(d)*). Once the pEF-mCitrine gene is silenced by HDAC4, it can no longer bolster the overall rate of reactivation at the pRSV-mCherry gene (*k_a1_(d)* is missing in Figure 6C compared to 6D) and protect it against background silencing. This reasoning is also consistent with the observation that we see less background silencing of the pRSV promoter when it is closer to the pEF in the no spacer construct compared to the 5kb one in the absence of CR recruitment (Figure 1).

In summary, both silencing and reactivation rates are affected by the distance between two genes. Upon recruitment of KRAB, a CR associated with reader-writer feedback and spreading of silencing histone modifications, the silencing rate is maximal near its recruitment sites and decreases slowly with distance (Fig 6E). However, HDAC4 recruitment leads to histone deacetylation, which is not associated with reader-writer positive feedback, and therefore is expected to decrease quickly over distance (Fig 6E). We hypothesize that CRs that are associated with positive reader-writer feedback loops can directly affect nearby genes, while CRs without feedback can only indirectly affect neighboring genes by changing the state of promoters very close to the recruitment site, which can in turn influence farther genes.

Promoters can be thought of as regions that are associated with high rates of gene activation and can drive reactivation after silencing. In the absence of chromatin regulator recruitment, each gene drives its own activation and in addition drives the reactivation of the neighboring gene in a distance-dependent manner (Figure 6F, top). In CHO-K1 cells, the pEF is a stronger promoter than the pRSV, while in K562 their strengths are closer (Figure 6F bottom, Figure 3–figure supplement 3). However, when a gene is silent, we are led to assume that the promoter can only reactivate itself and has no effect on reactivation of the adjacent gene until it is active again (Figure 6G). This assumption is necessary to explain why the silencing of pRSV after HDAC4 recruitment does not depend on its distance to the upstream silenced pEF promoter, even though an active pEF can increase pRSV reactivation in a distance dependent manner.

Similar to promoters, the core insulators can be modeled as DNA regions that increase the rate of reactivation of nearby genes in a distance-dependent manner (Figure 6H). This action, combined with the promoter reactivation rates, can prevent background gene silencing (Figure 6D) and drive reactivation after targeted CR-mediated silencing (Figure 6B&C). For example, because insulators increase reactivation rates, they can fight background silencing of the pRSV-mCherry both upon HDAC4 recruitment (Figure 6C), and under conditions without dox (Figure 6D). In K562, where the rate associated with pRSV activation is higher than in CHO-K1, the core insulators can bring the overall activation rate above background silencing when they are closer to pRSV. However, because the activation rates associated with insulators are small compared to KRAB-mediated silencing rates, they cannot insulate well against KRAB.

Thinking of promoters and core insulators as elements associated with higher reactivation rates also explains the distance-dependent gene reactivation after release of the chromatin regulators. When dox is removed after both genes are silenced, the reactivation dynamics are determined by the strength of the promoters and presence of insulators. In CHO-K1 cells, the stronger pEF gene and presence of any insulators leads to the reactivation of the mCitrine gene, which then can act at a distance to help reactivate the weaker pRSV-mCherry gene (Figure 6H, left). This leads to insulated reporters having the fastest reactivation and also explains the order of gene reactivation in the NS and 5kb reporters. In K562, where the strength of the pEF and pRSV promoters are more balanced, either gene can reactivate first and along with insulators drive reactivation of both genes (Figure 6H, right).

## Discussion

Characterizing how transcriptional silencing mediated by CRs affects neighboring genes is important for understanding interactions between adjacent genes in the genome and in synthetic circuits, as well as for developing safe gene therapy and epigenetic editing applications. To tease out the rules associated with these interactions, we engineered a series of synthetic reporters with different configurations, changing the distance between genes and testing different insulator arrangements in two mammalian cell lines. These reporters allowed us to monitor the extent and dynamics of spreading of silencing during recruitment of a chromatin regulator, as well as reactivation patterns and dynamics after its release. Using time-lapse microscopy and flow cytometry, we found that transcriptional silencing of a gene following recruitment of either KRAB or HDAC4 can affect a downstream gene even when separated by 5kb of distance or different insulator arrangements. KRAB silencing, associated with both histone deacetylation and H3K9 methylation, spreads with a delay that is very short (hours), increases with the distance between the genes, and is not affected by insulators. HDAC4 spreading, associated with histone deacetylation and H3K27me3, is slower (days) and does not depend on the distance between the genes. HDAC4-mediated silencing can also spread past insulators, though insulators seem to have a stronger effect on it, even blocking it in the DC configuration.

KRAB was previously shown to lead to spreading of both gene silencing and histone methylation when recruited near a gene as a fusion to either TetR^19, 20^ or to programmable DNA binding domains, such as dCas9^41^ or TALENs^17^. However, the extent of reported spreading varied in different contexts between 5kb to 200kb. Our results suggest that spreading of KRAB-mediated silencing is a dynamic process that depends on the time of recruitment as well as the level of activation at the target locus, tying together these apparently disparate results. For example, when KRAB is recruited for a short period at a highly active region such as the hemoglobin locus in K562, the extent of spreading of methylation is very small (<1 kb)^41^. In contrast, when KRAB is recruited for a long period (41 days) at a site with moderate acetylation, silencing can spread over 200kb^20^. Similarly, recruitment of HP1-alpha, a CR in the same silencing pathway as KRAB, has been shown to lead to silencing and spreading of H3K9me3 across ∼10kb over time ^42^.

The fact that KRAB-mediated silencing can spread to neighboring genes quickly raises two concerns for synthetic gene control and epigenetic editing. First, in mammalian genetic circuits where multiple genes need to be integrated in the same cell, genes controlled by KRAB need to be placed far away from other genes to avoid unwanted interference and feedback. Second, when targeting endogenous genes with KRAB (as in CRISPRi), the possibility of silencing spreading beyond the target gene should be considered.

In contrast to KRAB, silencing by HDAC4 (and in general by HDACs) is not traditionally associated with spreading of heterochromatin in mammalian cells, so we were initially surprised to observe that silencing mediated by HDAC4 can affect neighboring genes. In general, spreading of heterochromatin-mediated silencing is commonly associated with reader-writer feedback, which has been shown to contribute to heterochromatin spreading in theoretical models^43–46^ and synthetic experimental systems^47^. HDAC4 is not directly associated with CRs that can bind deacetylated histone tails; therefore, we did not expect to see new histone modifications after silencing. However, the appearance of H3K27me3, a PRC2 modification known to have reader-writer feedback, after HDAC4 recruitment led us to believe that the silencing of pEF-mCitrine allowed background silencing of pRSV-mCherry. This indirect silencing scenario could also arise upon silencing of endogenous promoters (for example in development, aging, or synthetic gene control), where their silencing would allow natural silencer elements in the gene neighborhood to suddenly start working.

We were surprised to discover that insulators were inefficient at blocking the spreading of chromatin-mediated targeted silencing, since insulators are traditionally thought to prevent spreading of heterochromatin and are commonly used in synthetic biology constructs to protect them from heterochromatin encroachment. None of the insulator variants derived from the cHS4 region of the beta-globin locus prevented spreading of silencing after KRAB recruitment (Figure 4). However, insulators did reduce background silencing of our reporters in CHO-K1 (Figure S11), consistent with results from classical transgene insulator assays^34, 39^. We also showed that even a single insulator can stop transcriptional run-on (Figure S2), consistent with previous reports^40^. We can explain the function of insulators on spreading and background silencing by thinking of them as areas in an active chromatin state, that can for example recruit additional writers of acetylation^38^. We propose that active regions associated with high acetylation, such as insulators and promoters, cooperate with each other to inhibit deacetylation. According to this model, insulators fail when the gene reactivation driven by insulators is slower than the KRAB-induced repression and spreading of silencing. However, insulators can work against a much slower silencing process such as background silencing. In other words, at these length scales, the process of “insulation” is better thought of as a dynamic fight between activation and repression. Screening different insulators^48^ in a similar recruitment assay in the future could help go beyond the binary classification of sequences as insulators or non-insulators and actually quantify how well they perform against different mechanisms of silencing. This process would also help identify more reliable insulators for synthetic biology.

By monitoring gene reactivation dynamics after release of the CRs, we conclude that activation can also spread to nearby genes in a distance-dependent manner. However, which gene reactivates first depends not only on the distance between the genes, but also on their promoter size and strength and overall reactivation propensity of the locus. These results led us to modeling promoters and insulators as regions with increased rates of reactivation. In this model, acetylation from strong promoters, such as pEF, can spread via looping and positive feedback to the downstream weaker RSV promoter, as seen in CHO-K1 cells. This model can be used in other contexts in which spreading of active modifications is relevant, such as enhancer-promoter interactions, and activation of nearby endogenous genes after epigenetic editing.

Together, our experimental results and the model based on them have broad implications for understanding chromatin mediated gene regulation and building mammalian synthetic biology applications. They suggest that genes in close proximity (<5-10kb) can respond to signals in a coordinated manner, during both silencing and activation. In our model, promoters, enhancers, and insulators can be represented simply as regions with increased rates of reactivation. It would be interesting to extend the system to scenarios when these regulatory regions are farther apart in linear space but nevertheless close in 3D-space, as is the case of many promoters and their corresponding enhancers or silencers. In terms of synthetic biology applications, this gene coupling can be detrimental when we want to deliver compact circuits of genes that need to be controlled independently. However, the length-dependent time delay in gene response can also be used to build more sophisticated temporal population responses. Finally, this experimental and theoretical framework can serve as a starting point for measuring and modeling the effects of targeting various epigenetic editors at endogenous loci in order to guide the time necessary for efficient on-target effects without unwanted off-target spreading.

## Acknowledgements

We would like to thank the members of the Bintu lab for discussions and helpful feedback, as well as Mike V. Van for cloning some CR plasmids. We especially thank Stanley Qi’s Lab for use of the Cytoflex cytometer, Mitsuo Oshimura’s Lab for the MI-HAC and phiC31 integrase, and Brooks Taylor (Mary Teruel’s Lab) for use of his cell-tracking software. This work was supported by a BWF-CASI Award (L.B.), NIH-NIGMS R35M128947 (L.B.), NIH-NIGMS Training Grant in Biotechnology 5T32GM008412 (S.L.), Japan Society for the Promotion of Science Overseas Research Fellowship (T.F.), NIH-NIGMS T32 Molecular Pharmacology Training Grant 5T32GM113854 (J.S.), NSF GRFP, VPGE Stanford Graduate Fellowship and VPGE EDGE Fellowship (M.M.H.), and NIH T32 Training Grant 5T32GM007365-45 (A.M).

## Author Contributions

S.L. and L.B. designed the study. S.L., M.H.H. and L.B. generated the DNA constructs and cell lines. S.L. and M.H.H. performed the flow cytometry experiments and associated data analysis with contributions from M.M.H., C.H.L., A.M. and L.B.. C.H.L. performed the CUT&RUN and associated analysis with contributions from S.L.. RNA extractions and subsequent analysis was performed by S.L. and M.M.H.. S.L. collected time-lapse microscopy data for which analysis was performed by J.S.. Model for time evolution of fluorescence distribution was derived by M.H.H., with input from L.B, and fit to flow cytometry data by M.H.H. Kinetic model of gene states was developed by L.B., S.L. and T.F.. S.L. and L.B. wrote the manuscript with contributions from all authors.

## Data and materials availability

All data, code, and materials used in the analysis will be publicly available upon publication.

## Supplementary Materials

### Materials and Methods

#### Plasmid construction

The CHO-K1 PhiC31 reporters (Figure S1A) were assembled as follows: First, a PhiC31-Neo-5xTetO-pEF-H2B-mCitrine reporter construct was assembled using a backbone containing the PhiC31 attB site, a neomycin resistance gene, and a multiple cloning site ^27^. The elements of the reporter constructs were PCR amplified from the following sources: five Tet binding sites from the TRE-tight plasmid (Clontech), pEF from pEF/FRT/V5-Dest (Life Technologies), and H2B-citrine from pEV2-12xCSL-H2B-mCitrine ^49^. These components were first sequentially cloned into the pExchange1 backbone using standard molecular biology techniques. The entire TRE-pEF-H2B-mCitrine was then PCR-amplified and combined by Gibson assembly with the phiC31-Neomycin-MCS backbone cut by AvrII. This construct was designed such that after integration, the neomycin gene would be expressed from a PGK promoter situated upstream of the phiC31 site in the MI-HAC ^27^. The second fluorescent reporter was added by digesting the mCitrine only plasmid with NdeI and adding: the pRSV-H2B from R4-blast-pRSV-H2B-mTurquoise (gift from Teruel Lab), mCherry from pEx1-pEF-H2B-mCherry-T2A-rTetR-EED (Addgene #78101), and polyA from PhiC31-Neo-ins-5xTetO-pEF-H2B-Citrine-ins (Addgene #78099) using Gibson Assembly to generate the NS construct. The lambda spacers were amplified from lambda phage DNA (NEB, N3011) and inserted via Gibson Assembly after digesting the NS reporter plasmid with Bsmb1. Similarly, the full length cHS4 was amplified from PhiC31-Neo-ins-5xTetO-pEF-H2B-Citrine-ins (Addgene #78099) or the core insulator was amplified from PB CMV-MCS-EF1α-Puro PiggyBac vector backbone (System Biosciences #PB510B-1), and inserted into the NS reporter with Gibson assembly after digestion with BsmBI for insulators between mCitrine and mCherry, and BsiWI (restriction digestion site added by site directed mutagenesis) for the insulators upstream of TRE. The PhiC31 integrase was a gift from the Oshimura Lab ^27^.

The K562 AAVS1 reporter constructs (Figure S1B) were assembled as follows: First, a 9xTetO-pEF-H2B-Citrine reporter was cloned into a AAVS1 donor vector backbone (Addgene #22212) containing a promoter-less splice-acceptor upstream of a puromycin resistance gene and homology arms against the AAVS1 locus. The elements of the reporter were amplified from the following sources: the 9XTetO sites were ordered from IDT, and the pEF-H2B-citrine was PCR amplified from the PhiC31 construct. These components were cloned into the AAVS1 donor vector backbone using Gibson Assembly. The mCherry components, spacers and insulators were added to the mCitrine only base plasmid in the same manner as the phiC31 reporters (see above), except for insulators where plasmids were digested with BstBI and MluI-HF.

The Piggybac plasmids containing the rTetR-CR were assembled into the PB CMV-MCS-EF1α-Puro PiggyBac vector backbone (System Biosciences #PB510B-1), which was modified via Gibson Assembly to add H2B-rTetR-Zeo from pEx1-pEF-H2B-mCherry-T2A-rTetR-KRAB-Zeo (Addgene #78352), mIFP from pSLQ2837-1, and for K562 plasmids the pGK promoter from pSLQ2818 (the latter two gifted from Tony Gao & Stanley Qi, Stanford). The CRs used were: rat KRAB from pEx1-pEF-H2B-mCherry-T2A-rTetR-KRAB (Addgene #78348) for CHO-K1, human KRAB ZNF10 from pSLQ2815 CAG-Puro-WPRE_PGK-KRAB-tagBFP-dCas9 (gifted from Tony Gao & Stanley Qi, Stanford) for K562, and human HDAC4 from pEx1-pEF-H2B-mCherry-T2A-rTetR-HDAC4 (Addgene #78349) for both CHO-K1 and K562.

All plasmids used in this study will be deposited to Addgene.

#### Cell Line Construction

Reporter lines in CHO-K1 cells were created by integrating the reporter plasmids at the phiC31 integration site on the MI-HAC (human artificial chromosome, gifted by Oshimura Lab) by co-transfecting 750 ng of reporter plasmid with 250 ng of the phiC31 integrase using Lipofectamine 2000 (Invitrogen, 11668027). Cells were plated 24 hours before transfection and media change was performed 12 hours after transfection. Selection was started 48 to 72 hours after transfection with 600 ng/mL geneticin (Gibco, 10131027) for one to two weeks. CRs were integrated randomly with the Piggybac system (System Biosystems, PB210PA-1) by co-transfecting 750 ng of CR plasmid and 250 ng PiggyBac transposase with Lipofectamine 2000. Cells were selected with 400 ng/mL Zeocin starting 24 hours or later after transfection for one to two weeks. Single clones were isolated for each lambda reporter, and correct integration into the MI-HAC was verified by PCR of genomic DNA. Note that rat KRAB was used for CHO-K1cells and human KRAB ZFN10 was used for K562; both are driven by CMV promoter and have mIFP as a fluorescent marker followed by T2A before the rTetR-CR fusion.

Reporters were integrated in K562 cells at the AAVS1 safe harbor site using TALENs: AAVS1-TALEN-L (Addgene #35431) targeting 5′-TGTCCCCTCCACCCCACA-3′ and AAVS1-TALEN-R (Addgene #35432) targeting 5′-TTTCTGTCACCAATCCTG-3′ ^50^. Roughly 1.2M K562 cells mixed with 5000 ng of reporter plasmid and 1000 ng of each left and right TALENS in a nucleofection cuvette (Mirus Bio, 50121), were transfected by nucleofection (Lonza 2B Nucleofector) with program T-16. Selection was started 48 to 72 hours after transfection with 3 ug/mL Puromycin (Invivo Gen, ant-pr) for one to two weeks. We performed genomic PCR on K562 multiclonal populations to confirm the presence of proper integration at the AAVS1 site, using primers from ^51^. CRs were integrated randomly with the Piggybac system by nucleofecting 1000 ng of CR plasmid and 300 ng PiggyBac. Cells were selected with 400 ng/mL Zeocin starting 24 hours or later after transfection for one to two weeks.

Both CHO-K1 and K562 reporter cell lines with CRs were sorted (Sony SH800) for triple positive fluorescence: mCitrine and mCherry of the dual gene reporter and mIFP transcribed along with the CR.

#### Cell Culture Conditions

Cells were cultured at 37°C in a humidified incubator (Panasonic MCO-230AICUVL) with 5% CO2. CHO-K1 cells were grown in Alpha MEM Earle’s Salts media with 10% Tet Approved FBS (Takara Bio 631367, Lots # A16039 & #17033) and 1X Penicillin/Streptomycin/L-glutamine (Gibco 10378016). CHO-K1 cells were passaged by rinsing with Dulbecco’s Phosphate-Buffered Saline (DPBS, Gibco 14190250), and incubating at room temperature with 0.25% Trypsin (Gibco, 25200056). K562 cells were grown in RPMI 1640 medium (Gibco, 11875119) with 10% Tet Approved FBS (Takara Bio 631367, Lot #17033) and 1X Penicillin/Streptomycin/L-glutamine (Gibco 10378016). For long-term storage, cells were frozen in growth media with 10% DMSO (Sigma Aldrich, D2650) and placed at −80°C (for up to a month), and then transferred to liquid nitrogen for long term storage.

#### Flow Cytometry of Recruitment and Release Assays

During recruitment assays, doxycycline (Tocris Bioscience, 4090) was added to the media to a final concentration of 1000 ng/mL unless otherwise stated. Fresh dox diluted from frozen aliquots was added to the media before each cell passage during recruitment, every 2-3 days. Flow cytometry data was collected on two different flow cytometers due to machine access: CytoFLEX S (Beckman Coulter) and ZE5 Cell Analyzer (BioRad). Cells were filtered through 40 um strainers to remove clumps before flow, and 10,000-30,000 cells were collected for each time point. For CHO-K1 NS and 5kb clonal lines were used for replicates; for the remaining multiclonal constructs, data for each time course was collected from experiments started on different days for use as biological replicates. Data was analyzed using a Matlab program called EasyFlow (https://antebilab.github.io/easyflow/) and a Python-based package, Cytoflow (https://cytoflow.github.io/). All cells were gated for mIFP positive cells based on fitting the background fluorescence of wildtype cells to a sum of Gaussians, and setting the threshold 2 standard deviations away from the mean of the main peak. The percentages of cells with mCitrine and mCherry active (Figure 1) were determined using a manual threshold based on the no dox sample. To quantify the percent of cells with mCitrine or mCherry ON during reactivation (Figure 5&S13&S14), we fit the background fluorescence of wildtype cells in each fluorescent channel to a sum of Gaussians and set the ON threshold 3 standard deviations away from the mean of the main peak.

#### CUT&RUN Experiments

CUT&RUN for the K562 KRAB and CHO-K1 KRAB cell lines was performed according to the third version of the published protocol on protocols.io^52^. For each antibody condition, 250,000 K562 or CHO-K1 cells were harvested and incubated with 10uL of activated Concanavalin A beads. Cells were permeabilized with 0.05% digitonin and incubated at room temperature for two hours with a 1:100 dilution of one of the following antibodies: guinea pig anti-rabbit IgG H&L chain (Antibodies-Online; Cat. No.: ABIN101961); rabbit anti-H3ac (Active Motif; Cat. No.: 39139); rabbit anti-H3K4me3 (Active Motif; Cat. No.: 39159); rabbit anti-H3K9me3 (Abcam; Cat. No.: ab8898). Unbound antibody was washed out, and cells were incubated with 140ng/mL of a protein A and micrococcal nuclease fusion (pA-MNase; generously provided by the Henikoff lab). Unbound pA-MNase was washed out, and cells were resuspended in a low-salt buffer. Tubes were placed in a thermal block chilled to 0°C, and 10mM CaCl2 was added to trigger DNA cleavage by pA-MNase. After 5 minutes, the reaction was stopped by chelating calcium ions with the addition of 20mM EGTA, and tubes were incubated at 37°C for 30 minutes under physiological salt conditions to promote diffusion of fragmented chromatin into the supernatant. Fragmented chromatin in the supernatant was collected and purified using the DNA Clean & Concentrator-5 kit (Zymo; Cat. No.: D4004). CUT&RUN for the CHO-K1 HDAC4 cell lines was performed using the EpiCypher CUTANA ChIC/CUT&RUN Kit (Cat. No.: 14-1048) according to the User Manual Version 2.0 with the following conditions: an input of 500,000 CHO-K1 cells per antibody condition; a final digitonin concentration of 0.01%; an overnight antibody incubation at 4°C with 1:100 dilutions of the anti-H3Kac and anti-H3K9me3 antibodies described above along with a 1:50 dilution of rabbit anti-H3K27me3 (Cell Signaling Technologies, Cat. No: 9733S); and DNA purification with the provided spin columns. Dual-indexed sequencing libraries were prepared using the NEBNext Ultra II DNA Library Prep Kit for Illumina (New England Biolabs; Cat. No.: E7645L) with 12-14 cycles of PCR, and size selection was performed with either Agencourt AMPure XP (Beckman Coulter; Cat. No.: A63880) or SPRIselect (Beckman Coulter; Cat. No.: B23318) magnetic beads. Library concentrations were quantified with the Qubit 1X dsDNA HS Assay Kit (Invitrogen; Cat. No.: Q33231), and library fragment sizes were assessed with the High Sensitivity D1000 ScreenTape (Agilent; Cat. No.: 5067-5584) and High Sensitivity D1000 Reagent (Agilent; Cat. No.: 5067-5585) on an Agilent 4200 TapeStation System. Libraries were pooled for genome-wide sequencing.

#### CUT&RUN Sequencing, Data Processing, and Data Analysis

Genome-wide libraries were paired-end sequenced by the Stanford Center for Genomics and Personalized Medicine on a HiSeq 4000 with 2 x 101 cycles. A custom genome with our reporter sequence appended to the end of the hg19 human genome assembly was constructed with bowtie2-build. Paired-end alignment was performed with the following bowtie2 command: bowtie2 –local --very-sensitive-local --no-unal --no-mixed --no-discordant --phred33 -I 10 -X 700 -x {reference genome} -p 8 −1 {first mate of pair} −2 {second mate of pair} -S {output SAM file name}. Fragments that mapped completely within non-unique reporter elements (i.e. pEF, H2B, PGK, AmpR, or SV40 polyA) were ambiguous and thus removed to avoid confounding. Samtools was used to convert SAM files to BAM files and to subsequently sort and index BAM files. Picard was used to mark and remove duplicates with the following command: java -jar {picard tool} MarkDuplicates -I {input sorted BAM file} -O {output deduplicated BAM file} -M {output metrics file} --REMOVE_DUPLICATES true. Reads were normalized to counts per million (CPM) and bedgraph files were generated with the following bamCoverage command: bamCoverage --bam {input deduplicated BAM file} -o {output bedgraph file} --outFileFormat bedgraph --extendReads --centerReads --binSize 10 --normalizeUsing CPM. \CUT&RUN data were visualized via the Broad Institute and UC San Diego Integrative Genomics Viewer. Additional data analyses and visualization were performed with custom scripts in Python.

ChromaBlocks analysis^53^ was performed in R using the Repitools package. Normalized bedgraph files were read in as dataframes and converted to GRangesList objects with the annoDF2GR() and GRangesList() methods. GRangesList objects for H3K9me3, H3Kac, or H3K4me3 samples were specified for the rs.ip argument within the ChromaBlocks() method, and the respective dox-treated or untreated IgG sample was specified as the rs.input argument. ChromaBlocks() was called with the ‘large’ preset for H3K9me3 and H3Kac and with the ‘small’ preset for H3K4me3 to identify regions of enrichment. Start coordinates and widths of enriched regions were used to construct bed files for visualization in IGV and for signal integration to compute the log2 fold-change in Figure S1E.

#### qPCR and PCR on cDNA

Each cell line was treated with dox (1ug/mL) for the indicated number of days before harvesting. RNA was harvested using RNeasy Mini Kit (Qiagen, 74106) using the on-column RNase-Free DNase (Qiagen, 79254). cDNA was synthesized using the iScript Reverse Transcription Supermix (Biorad, 1708840) for qPCR experiments and qScript cDNA SuperMix (VWR, 101414-106) for PCR experiments. qPCR on cDNA was performed using SsoFast EvaGreen Supermix on a CFX Connect Real-Time PCR System (both from Bio-Rad Laboratories). For qPCR primer sequences, see Table S1. The values are reported as delta Ct to Beta-actin. PCR on cDNA was performed using Q5 Hot Start High-Fidelity 2X Master Mix (NEB, M0494L) on a C1000 Touch Thermal Cycler from Bio-Rad. PCR products were run on a 1-1.5% TAE agarose gel at room temperature and imaged using Biorad Gel Doc EZ Imager. For PCR primer sequences, see Table S1.

#### Time-lapse Microscopy

Fluorescent time-lapse imaging was conducted using a DMi8 inverted epifluorescence microscope (Leica, Germany) with a 20X Plan Apo 0.8 NA objective and a Leica DFC9000 GT CMOS Camera using 2×2 binning. The microscope was enclosed in a cage incubator (Okolab Bold Line, custom for DMi8) held at 37°C, and samples were placed in a stage top chamber (Okolab H301-K-FRAME) also held at 37°C, 5% CO2 (Praxair Cat. BI NICD5O6B-K), and with humidity control. CHO-K1 cells were seeded 24 hours prior to the start of movie acquisition on optically clear 96-well µ-Plates (Ibidi Cat. 89626). Border wells were filled with PBS (Thermo Fisher Scientific Cat. 10010023) and plates were sealed with Breath-Easy gas-permeable membranes (Diversified Biotech Cat. BEM-1) to reduce evaporation and maintain humidity during image acquisition. Cells were imaged in alpha MEM media with no phenol red (Thermo Fisher, Cat. 41061029) with 10% Tet-Free FBS (Takara Cat. 631106; Lot #A17033), and 1% Penicillin/Streptomycin/Glutamine (Fisher Scientific Cat. 10378-016) to reduce background fluorescence. Images were acquired every 20 minutes in three different fluorescent channels: YFP, RFP and CY5, corresponding to the mCitrine, mCherry and mIFP fluorophores, respectively. The light source was a Sola Light Engine, and a Lumencor control pod was used to attenuate the light to <1mW range. The FIM was set to 10% and Lumencore was set to 5%. Exposure times were 150 ms, 200 ms and 100 ms for the CY5, RFP and YFP channels, respectively (an additional trigger cable between the light source and microscope was necessary to achieve precise exposure times). One to two time-lapse movies were analyzed for each cell line, where each movie consisted of cells in dozens of different wells, and in each well 3-5 non-overlapping sites were imaged. Cell culture media was changed every 24 hours to replenish doxycycline and remove toxic reactive oxygen species which may have formed due to prolonged imaging. LasX software was used to control the microscope in the Mark and Find module to image up to 200 positions in total and AFC on demand mode adjusted focus for every cycle and position of imaging.

#### Analysis of Time-lapse Movies

Analysis of all time lapse movies was conducted in MATLAB R2016a (MathWorks) unless otherwise stated. Fluorescent imaging data were obtained by automated image segmentation and single-cell tracking using the MACKtrack package (https://github.com/brookstaylorjr/MACKtrack). Nuclei were segmented and tracked using H2B-mIFP signal or H2B-mCherry signal (on a case by case basis based on expression level) and single-cell intensities for integrated nuclear intensity of mCitrine, mCherry and mIFP channels were measured at each frame. Tracked single-cell output traces were then filtered in a semi-automated fashion to remove incomplete or mistracked cells which were defined as traces exhibiting numerous aberrant drops in fluorescence intensity inconsistent with cell divisions. Additionally, traces consisting of background silenced cells with low mCitrine expression prior to the 24 hour time point were omitted from analysis. In order to call transcriptional silencing events, the gradient of the cumulative intensity traces for mCherry and mCitrine were used. To generate cumulative traces, mitotic events were first computationally identified based on periodic dilution of the stable H2B-fluorophore signal by approximately 50% upon cell division. Using these points as reference, raw cumulative traces for mCherry and mCitrine were generated by computationally re-adding the lost fluorescence due to cell division events and adding these values to all subsequent time points after a division event to “stitch together” intensity values across divisions (see Figure S2A). The raw stitched traces were smoothed to obtain smoothed stitched traces by applying: first a moving average filter (“smoothrows” function, Matlab) with a window size of 2 frames, then a “smoothingspline” (“fit” function, Matlab) with the smoothing parameter set to 1.0e-04, and finally a median filter (“medfilt1” function, Matlab) with a window-size of 3 frames. The gradient of these smoothed stitched traces were used to call silencing when it fell below a threshold. mCitrine or mCherry silencing was called in individual cells at the first frame where the gradient of the smooth cumulative trace fell below a threshold (25% for KRAB and 30% for HDAC4) of its initial value (taken to be the maximum gradient of the same trace between frames 50-70 prior to dox addition). Furthermore, cells must have stably remained below that threshold through at least one more cell division to be considered silenced. To account for the spike in mCherry signal following mCitrine silencing in certain cells (Figure S4B), the gradient threshold was called in these instances relative to a 10% drop from its absolute maxima which represented the spike directly prior to transcriptional silencing. Single-cell silencing delays were calculated by subtracting the frame number at which silencing was called in mCitrine from the corresponding silencing frame number of mCherry in the same cell and converting to hours.

#### Statistical Analysis of Movie Delay Times

Welch’s unequal variances T-test was performed on GraphPad Prism 8.4 on the calculated time delays from movies to determine significance (p-values) between chromatin-regulator spreading rates between NS and 5kb reporters.

#### Model for the Time Evolution of Fluorescence Distributions

We have developed a model that describes the time dependent evolution of the fluorescence distributions of our two fluorescent reporters after the recruitment of CRs. This model has two components: a deterministic component describing the decay of fluorescence due to mRNA and protein degradation and dilution, and a probabilistic component taking into account that cells can stochastically transition between the active and silent states at different times. In the deterministic component, we will derive equations that describe the fluorescence of cells in either the active or silent state, beginning with the simpler upstream *mCitrine* gene, followed by the downstream *mCherry* gene. The deterministic equations are then used in the probabilistic component to derive a probability density function that describes flow cytometry data. By deriving this model and fitting it to our data, we estimate the silencing parameters of our system such as the time delay between reporter silencing and the magnitude of transcriptional interference.

##### Deterministic Component with mRNA and Protein Degradation and Dilution

In this component of the model, we derive deterministic equations for the fluorescence of cells in the active and silent state. First, we will derive these equations for the simpler upstream *mCitrine* gene. We begin at the level of transcription where *M* represents the concentration of reporter mRNA. The recruitment of the CR begins at *t=0* with the addition of doxycycline, but transcriptional silencing of *mCitrine* does not occur until some time later, *t_s,c_*. By assuming a constant mRNA production rate *α_m_*, and a constant coefficient of mRNA degradation and dilution, *β_m_*, we can write a piecewise differential equation for mRNA concentration:

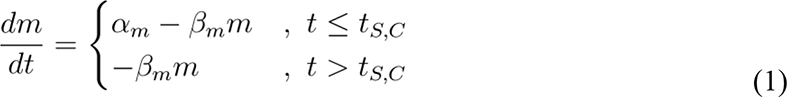

This differential equation describes transcription of the upstream *mCitrine* gene. We assume a steady-state initial condition and continuity among the pieces to arrive at the following solution for *mCitrine* mRNA as a function of time:

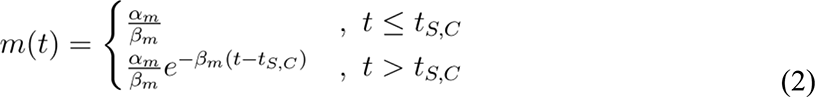

Next, we move on to the level of translation, where mRNA is translated into protein. The protein concentration is represented by *ρ*. Here we assume a constant coefficient of protein production, *α_p_*, and a constant coefficient of protein degradation and dilution, *β_p_*. These assumptions allow us to write the following general differential equation to represent translation:

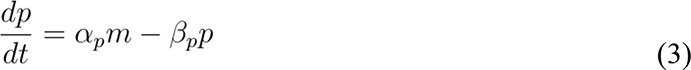

Substituting our solution for *m(t)* into the equation above and combining *α_p_* and *α_m_* into a single integrated production parameter such that *α = α_p_α_m_*, we arrive at the following piecewise differential equation for the concentration of the mCitrine protein over time:

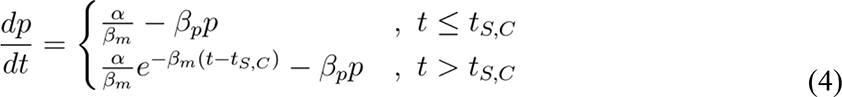

Again, we assume a steady-state initial condition and continuity among the pieces to arrive at the following solution for mCitrine concentration:

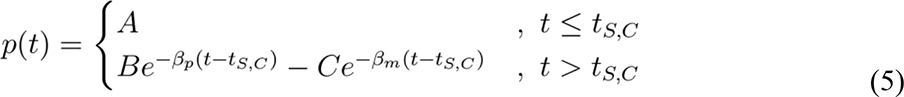

Where

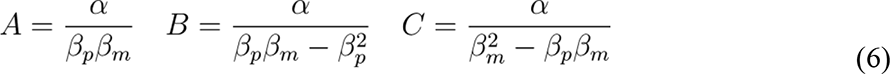

Now that we have derived equations for the concentration of protein over time, we need to convert from concentration to fluorescence, represented by *x* in our model. We assume that fluorescence increases proportionally to concentration after the addition of some background fluorescence *x_B_*. The proportionality constant is absorbed into the production parameter, α. Therefore, fluorescence can simply be expressed as protein concentration with the addition of the background fluorescence in Arbitrary Fluorescence Units (AFU):

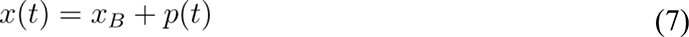

Using this relationship and the solution to *p(t)*, we define two separate equations for mCitrine fluorescence, one for the active state where *t* ≤ *t_S,C_* and one for the silent state where *t* > *t_S,C_*:

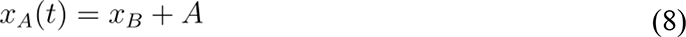

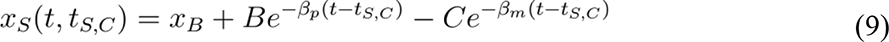

These two equations describe the fluorescence for the upstream *mCitrine* gene, but they are too simple to describe silencing at the downstream *mCherry* gene, as we often observe an increase in mCherry after the recruitment of the CR. We hypothesize that this upwards spike in mCherry production is due to the loss of transcriptional interference upon silencing of the upstream *mCitrine* gene. Therefore, we modified the differential equation for mRNA with the addition of a new term, *γ*, which represents the factor by which mRNA production is reduced by transcriptional interference. This factor is removed after the average silencing time of the *mCitrine* gene, *t_S,C_*, causing a delayed spike in the mRNA concentration of *mCherry* before it silences and begins to degrade at its own *t_S,M_*:

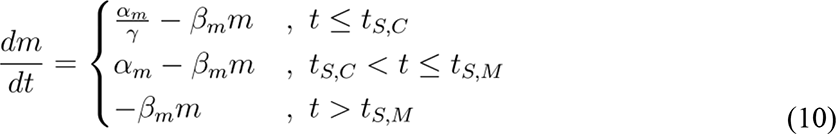

To solve this differential equation, we again assume a steady-state initial condition and continuity among the pieces. The solution for *mCherry* mRNA over time is:

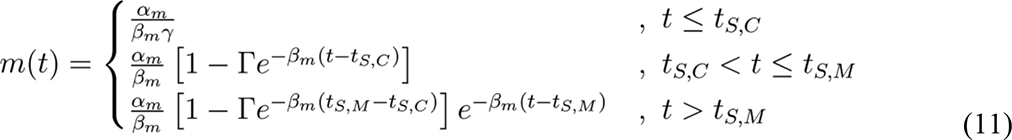

Where

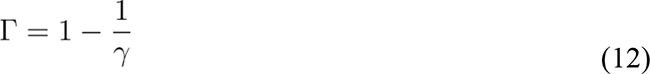

Substituting this solution for *m(t)* into the general translation equation and again combining α’s into an integrated production parameter α, we arrive at a piecewise differential equation for the concentration of mCherry over time:

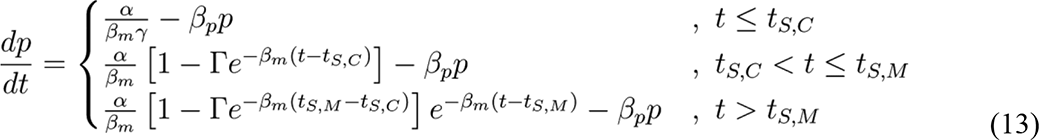

Again, we assume a steady-state initial condition and continuity among the pieces to arrive at the following solution for *mCherry* concentration over time:

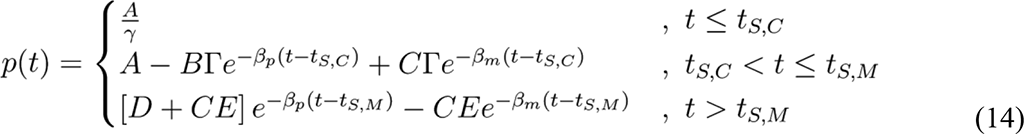

Where

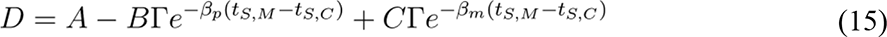

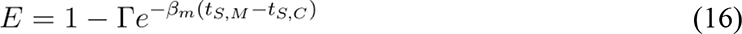

While the previous equation for *p(t)* assumes the more likely scenario that the upstream *mCitrine* gene silences before the downstream *mCherry* gene (*t_S,M_* > *t_S,C_*), we must also consider that *mCherry* may silence before *mCitrine* (*t_S,M_* ≤ *t_S,C_*). In this scenario, there is no spike of mCherry expression because the *mCherry* promoter silences without the loss of transcriptional interference concomitant with *mCitrine* silencing. Therefore, the following solution for mCherry concentration over time when mCherry silences first is the same as mCitrine, only divided by *γ*:

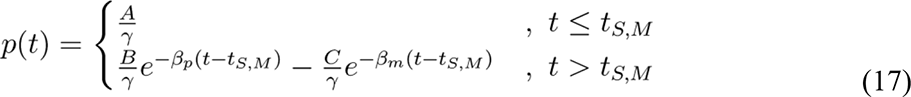

Now that we have defined two equations of mCherry concentration, we again combine these equations with the fluorescence equation and split them such that there is one equation for the active state where *t* ≤ *t_S,M_* and one equation for the silent state where *t* > *t_S,M_*. The active state fluorescence combines the first two parts of equation 14 with the first part of equation 17 while the silent state fluorescence combines the third part of equation 14 with the second part of equation 17. These combinations reflect that in either the active or silent state, either reporter can silence first. The fluorescence equations for mCherry are:

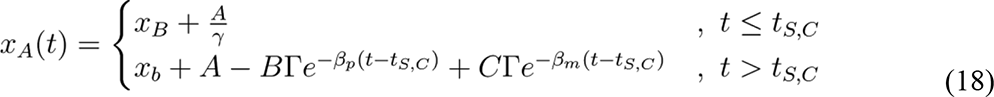

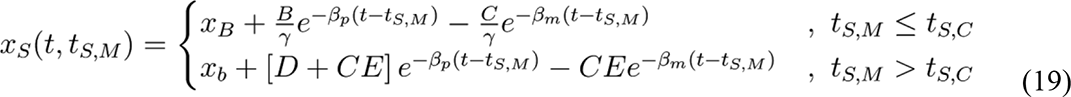

These two equations describe the downstream mCherry fluorescence at any given point in time and with any given *t_S,C_* and *t_S,M_*. They will be useful in the next section of the model. The constant parameters of the model thus far are as follows:

- α - the integrated fluorescence production parameter in AFU/day^2^
- *β_p_* - the protein degradation and dilution coefficient in days^-1^
- *β_m_* - the mRNA degradation and dilution coefficient in days^-1^
- *x_B_* - the background fluorescence in AFU
- γ - the interference factor (only applicable for mCherry)

##### Probabilistic Component with Stochastic Transcriptional Silencing

In the probabilistic component of the model, we include a stochastic transition between active and silent states in the derivation of a probability density function of fluorescence *x* at time *t*, *f(x,t)*. The distribution evolves over time and describes the flow cytometry data that we collect over the course of our silencing experiments. This probability density function of fluorescence is a weighted sum of three independent probability density functions, one for each of three possible cell states: background silent cells, *f_B_(x)*; active cells, *f_A_(x, t)*; and silent cells, *f_S_(x, t)*;. The time dependence in active and silent cells reflects the time dependence in the deterministic equations describing fluorescence in the active and silent states. Furthermore, because we have observed that the transition from the active to the silent state is a stochastic process with variability between cells, ^8^ we incorporate a time dependence into the weights of active and silent cells, *P_A_(t)* and *P_S_(t)* respectively. The probability density function can hence be written:

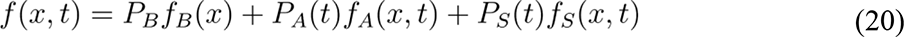

To consider all cells and for *f(x,t)* to meet the definition of a probability density function, the weights of each state, represented by P’s, must sum to 1. For simplicity, we assume that the fraction of background silent cells, *P_B_*, does not significantly change over the silencing time course. Therefore, *P_B_* does not have a time dependence in the following relationship:

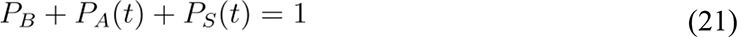

To determine the fraction of active cells and silent cells at any point in time, we consider that a cell transitions from an active state to a silent state after a time of *t_S_*. We add stochasticity to this transition by allowing cells to transition at different times and defining random variable *T_S_*. We choose a gamma distribution for *T_S_* as this distribution is defined to be strictly greater than zero and can assume various shapes between an exponential distribution with *K* = 1 and a normal distribution as *k* approaches infinity. In the shape-rate parameterization of the gamma distribution, the rate, typically represented as θ, can be expressed by dividing the mean, *τ_S_*, by the shape *k*. Therefore, we attain the following definition and probability density function for *T_S_*:

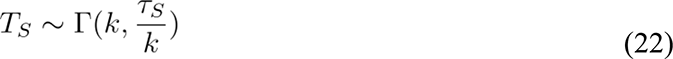

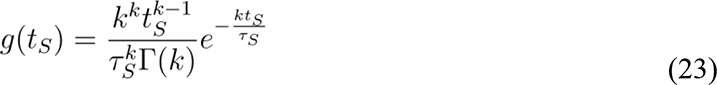

Using this definition, we will attain solutions for *P_A_(t)* and *P_S_(t)* by first integrating over *T_S_*’s probability density function from 0 to *t* to yield the following cumulative density function:

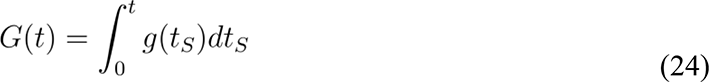

This cumulative density function represents the fraction of active cells that have transitioned into the silent state until and must be multiplied by (1 - *P_B_*) to yield the overall weight of silent cells:

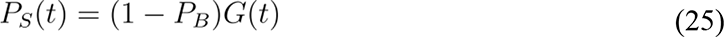

To get the weight of active cells, we simply subtract the cumulative density function from 1 to find the fraction of active cells that have not silenced until before multiplication by (1 - *P_B_*):

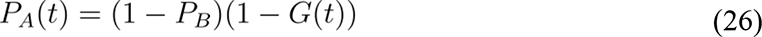

With analytical expressions for each weight, we now consider each independent probability density function of fluorescence for each of the three cell states. First, the probability density function for the fluorescence of background silent cells does not have a time dependence and is log-normal with a median of the background fluorescence, *x_B_*, and a shape parameter, or log-space standard deviation, σ:

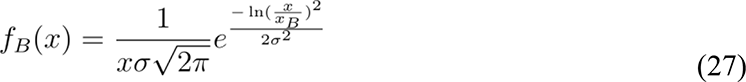

Next, the probability density function for the fluorescence of active cells is also log-normal, but the median of this distribution is given by *x_A_(t)* derived in the previous section. For simplicity, we assume that the shape parameter of this distribution, σ, is the same as the background distribution:

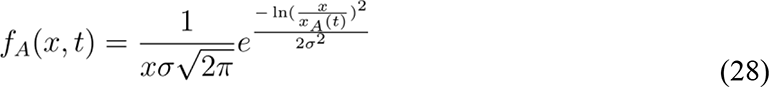

Finally, the probability density function for the fluorescence of silent cells is a convolution of *T_S_*’s gamma distribution for silencing and a log-normal distribution with a median of *x_S_(t, t_S_)*. The convolution occurs over *t_S_* from time 0 to *t*. This step can be thought of as a weighted sum of log-normal distributions with different medians based on when silencing occured. Because the weights only sum to the probability of silencing up to *t*, the integral must be normalized by division by *G*(*t*). This normalization ensures that *f_S_(x, t)* integrates to one:

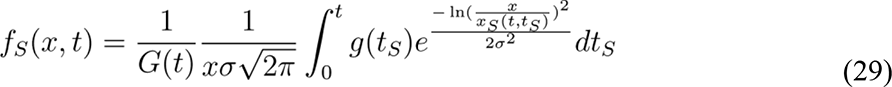

Again, we used the same shape parameter, σ, so that when the fluorescence decays to background levels, the distribution for fluorescence of background silent cells matches the distribution for silent cells:

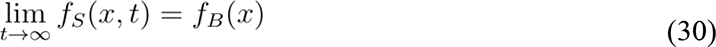

We have now defined the weights and probability density functions for each of the three possible states in terms of some new parameters and the equations derived in the deterministic component, which concludes the probabilistic component of the derivation. The additional constant parameters of the model are:

- *P_B_* - the fraction of background silent cells
- *k*-the shape parameter of *T_S_*’s gamma distribution
- *τ_S_* - the mean silencing time from of *T_S_*’s gamma distribution in days
- σ - the shape parameter for all log-normal fluorescence distributions

##### Fitting Method

Fitting our probabilistic model to the flow cytometry data allows us to estimate the relevant silencing parameters. Of particular interest are the interference parameter, γ, and the mean silencing times for *mCitrine* and *mCherry*, *μ_S,C_* and *μ_S,M_*, as well as the difference between the two, or Δ *μS*. Our model fitting was conducted in Python 3 Jupyter notebooks.

First, we defined functions F_o(t,params*) and F_s(t,t_l,params*) to calculate *x_A_(t)* and *x_S_(t, t_S_)* from the deterministic portion of the derivation respectively. Within these functions, we set the mRNA half-life *β_m_*) to four hours as was previously determined experimentally ^8^. Then we defined f(x,t) to calculate the probability distribution derived in the probabilistic portion of the derivation. Although the derivation describes the distribution on a linear scale, f(x,t) calculates the distribution on a log_10_ scale as this scale is more commonly used to visualize and work with flow cytometry data.

With functions defined to calculate the model solution given a set of parameters, we next defined an error function to calculate error between the data and the model. To calculate this error, the fluorescence values are binned into 100 equally spaced bins for each timepoint in the data. The probability densities of each bin are calculated. Then, we evaluate the model with the given parameters at the midpoint of each bin. The error is calculated by subtracting the bin densities from the solution to the model’s probability density function. The error arrays from each timepoint are appended to generate a single array of errors for the whole time course.

The mCitrine and mCherry reporters were each fit separately and had differences in the parameters used. Therefore, we used the general error function described above to define two separate error functions for mCitrine and mCherry. Since we never saw a spike in mCitrine expression, the interference parameter, *γ*, was intentionally set to one and the spike delay, *t_S,C_* in the deterministic derivation for mCherry, was set to zero. Fixing these parameters eliminates the reporter spike and allows us to use the same set of equations to describe both mCitrine and mCherry. Additionally, we set the shape parameter of *T_S_*’s gamma distribution for mCitrine to one. This constrains the gamma distribution to an exponential distribution, which is more suitable for mCitrine because this reporter silences quickly and reliably upon induction (Figure 2, time-lapse movie results).

The mCherry fit was conducted after the mCitrine fit as some of the parameters are carried from mCitrine to mCherry. One of these parameters is the protein dilution *β_p_*, which can be more reliably calculated in the mCitrine fit due to higher dynamic range and more uniform silencing. Since both the mCitrine and mCherry reporters are fused to H2B, they are stable and do not degrade significantly over the course of the experiment. Therefore, *β_p_* is governed by dilution due to cell division and is the same for both mCitrine and mCherry. In addition, *Τ_S,C_* was calculated in the mCitrine fit and then used as *t_S,C_* in the mCherry deterministic equations.

There is one additional consideration for the mCherry fits. Due to fast mCherry silencing by KRAB, it was difficult estimate both the interference factor, *γ*, and the mean silencing delay, *μ_S,M_*. Therefore, we used the *γ*’s estimated from the slower HDAC4 silencing time courses for each reporter when estimating the parameters in the KRAB silencing data. We believe this is acceptable because transcriptional interference is a property of the reporter and not the CR. The exception is that for the K562 insulator lines we did not fit the model to the HDAC4 data because we do not see complete silencing of mCherry. For these HDAC4 lines we estimated *γ* by taking the ratio of the maximum mode of the fluorescence in the time course to the mode on day zero and used this estimated *γ* for the KRAB fits.

One final adjustment to the model was made to fit non-saturating dox concentrations (100ng/mL and 200ng/mL). When dox concentration was low, we did not observe silencing of mCitrine in all cells as was the case for 1000ng/ml, and accordingly did not observe subsequent spreading of silencing to mCherry in those cells. To account for only a fraction of cells being silenced, an additional parameter was added to the model to represent cells that are always active, *P_aa_*. These cells do not silence, and their fluorescence distribution is fixed at active-state levels. Consequently, only 1-*P_aa_* are involved in silencing and are subject to the components described by equation (21).

To estimate the unconstrained parameters for each reporter, we fit the model to the data using a least-squares approach, scipy.optimize.least_squares(). This function solves for the parameters that minimize the sum of the squares of the error array that we defined. Initial guesses for each parameter were reasonably estimated by manual inspection of the data where possible. Bounds for each parameter were set generously as to avoid bounding of the solution where possible. The fits were performed independently for each replicate or single clone and took several hours for each cell line. However, the resulting fits were satisfactory by visual inspection, and the parameters were in accordance with those found in time course movie analysis. The 90% confidence interval (CI) for the parameters extracted from fitting daily flow cytometry data to the model, was estimated using a t-distribution with t-score 2.92 and sample standard deviation from all replicates.

#### Supplementary Text

##### Transcriptional Interference

We noticed an increase in mCherry expression after 5 days of HDAC4 recruitment in K562 with the NS reporter (Figure 1E). We wondered if this spike in mCherry fluorescence was due to ceasing of transcriptional interference on the pRSV-mCherry from the upstream pEF promoter. Transcriptional interference can occur through promoter occlusion as has been previously measured ^29, 54, 55^. We hypothesized that if transcription termination failed at the mCitrine polyA, Pol II would run on into the RSV promoter and hinder transcription initiation of the mCherry gene (Figure S2A). l Upon transcriptional silencing of the upstream pEF promoter, regular transcription initiation at pRSV can resume, leading to the observed temporary increase in mCherry expression prior to chromatin mediated silencing of pRSV-mCherry (Fig S2B). To test if any transcripts span from the mCitrine gene body across pRSV into the mCherry gene body, we performed PCR on cDNA from K562 cells after different durations of HDAC4 recruitment. This revealed that indeed, in cells where pEF has not yet been silenced by HDAC4, there is transcriptional run-on from pEF over pRSV, and that run-on does not occur in cells where mCherry is increased (Fig S2B). This suggests that the observed increase in mCherry production is due to pEF silencing and transcriptional interference no longer occurring. We do not see this effect with KRAB because silencing spreads more quickly than for HDAC4, and does not leave sufficient time for transcription to initiate at pRSV before both genes are silenced.

To experimentally measure the dynamics of transcriptional interference, we performed RT-qPCR against different elements of the reporter after 0, 1 and 5 days of HDAC4 recruitment in CHO-K1 cells with the 5kb reporter (Figure S2C-D). Our data confirmed transcription of the 5kb lambda region, and revealed that mCitrine and lambda mRNA levels both decreased after one day of HDAC4 recruitment. This further suggests that transcription runs on from mCitrine through the 5kb lambda into mCherry, preventing transcription initiation at pRSV. Interestingly, mCherry mRNA abundance did not change between 0 and 1 day of HDAC4 recruitment, even though the mCherry protein level significantly increased after 1 day of HDAC4 recruitment. Since it is not possible to distinguish the initiation point of transcripts by RT-qPCR, the measured mCherry mRNA abundance comes from a combination of transcripts initiated at pRSV and transcripts initiated at pEF that run through mCherry. It should only be possible to translate mCherry from transcripts initiated at pRSV since the mCherry in the run-on transcript would not be in the correct reading frame. Therefore,the observed increase in mCherry protein abundance may result from an increase in the fraction of translatable mCherry mRNA as pEF is silenced. Further experiments would be needed to confirm this.

We noticed that in the absence of dox pRSV-mCherry expression for the SH reporter is much higher than for the 5kb lambda reporter in CHO-K1 (Figure S12). We also do not observe an increase in mCherry expression upon HDAC4 recruitment in CHO-K1 cells with insulators(Figure S2E), unlike in the 5kb lambda reporter (Figure S2D). We performed PCR on cDNA to test if any transcripts span from the mCitrine gene body across the cHS4 insulator and pRSV into the mCherry gene body at different durations of HDAC4 recruitment in CHO-K1. The run-on transcript was not present at day 0 or after 1 or 5 days of HDAC4 recruitment in the SH insulator by PCR (Figure S2F). This suggests that the single cHS4 insulator, and possibly the cHS4 core sequence, can terminate transcription. This effect is in agreement with previous studies which have shown that insulator sequences can enhance gene expression and reduce promoter interference^40^.

##### Background Silencing

The extent to which background silencing is mitigated is influenced by the insulator configuration, cell type, and the CR that is over-expressed. By monitoring mCitrine and mCherry expression over time without CR recruitment, we are able to quantify the fraction of cells that spontaneously silence either mCitrine or mCherry (Figure S11). We calculated the rate of background silencing over time for each reporter by performing a least squares regression to the fraction of background silenced cells for each clone or replicate (Figure S11). For both KRAB and HDAC4 in CHO-K1 cells, all the insulator configurations reduced background silencing of the mCitrine and mCherry genes in comparison to the NS and 5kb reporters. By contrast, the insulators increase the rate of background silencing in some configurations in K562 cells, and this effect is dependent on the CR present. With KRAB, all the insulator configurations lead to slightly higher background silencing than the NS and 5kb reporters, and the level of silencing is similar for mCitrine and mCherry. With HDAC4, the levels of background silencing differ between mCitrine and mCherry, and the SH and DH insulators have strikingly high levels of mCherry background silencing. In only switching either the CR or cell type in a controlled system, we show that insulators have vastly different effects on neighboring genes.

---

**Figure 1 - figure supplement 1.**
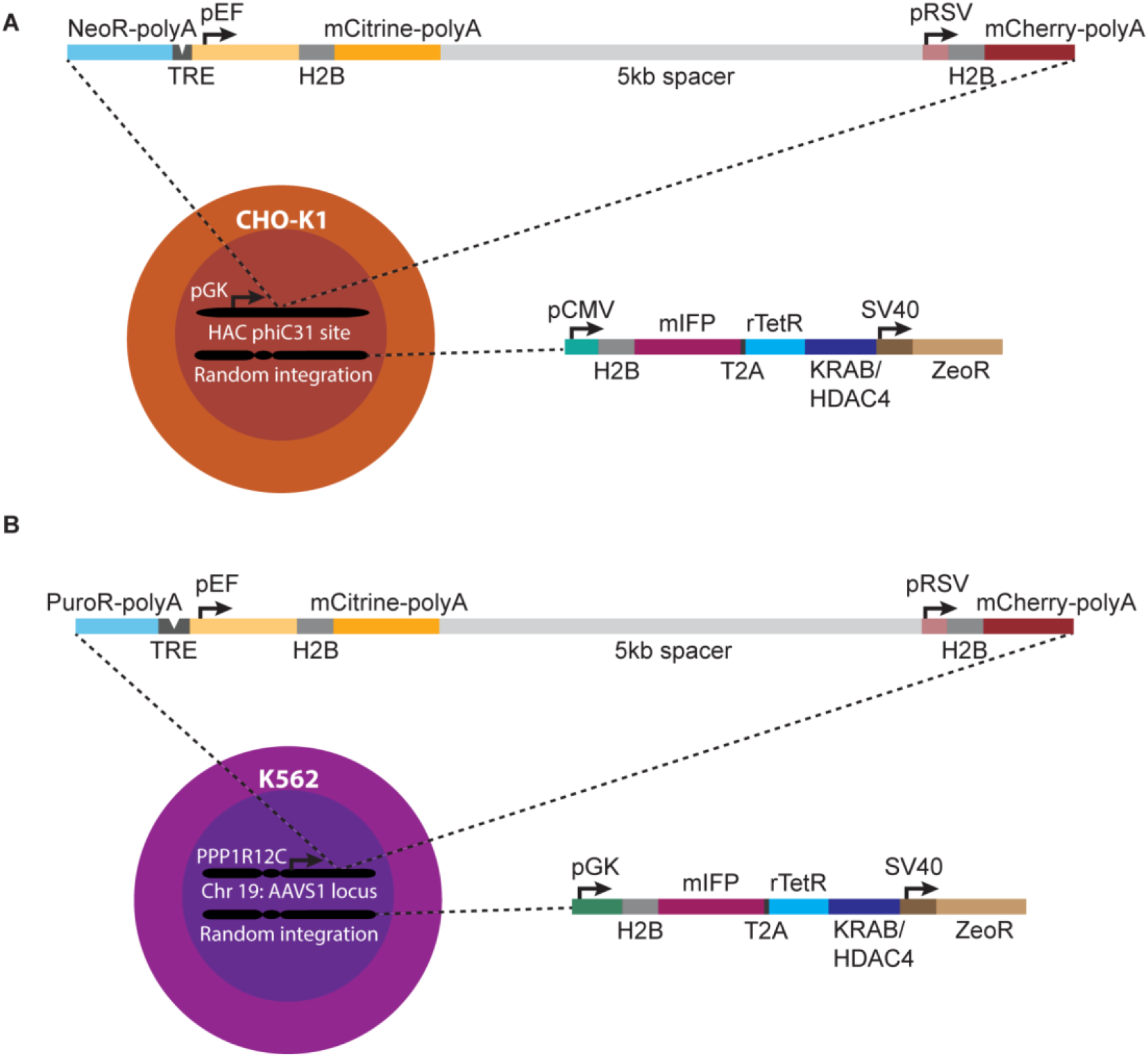
Reporter constructs used in different cell lines for analyzing spreading of transcriptional changes. **(A)** Dual mCitrine-mCherry reporter (top) is integrated in CHO-K1 cells in the MI-HAC ^56^ at the phiC31 integrase site. Upon integration, the neomycin resistance gene (NeoR) starts being expressed from the pGK promoter in the MI-HAC. **(B)** Dual mCitrine-mCherry reporter is integrated in K562 cells by TALENS at the AAVS1 locus. Upon integration, the puromycin resistance gene (PuroR) is driven by the promoter for the PPP1R12C gene ^28^. Both mCitrine and mCherry genes have SV40 polyA tails which should aid in transcriptional termination of each gene. NeoR in CHO-K1and PuroR in K562 both have a BGH polyA. The TRE in CHO-K1 has 5 TetO binding sites, while the TRE in K562 has 9 TetO binding sites. In both cell types, each fluorescent protein is fused to H2B which stabilizes the fluorophore and localizes it to the nucleus, allowing better tracking of single cells in time-lapse microscopy. Chromatin regulators fusions of rTetR to either KRAB or HDAC4 are expressed via random integration by piggyBac transposase, the approximate expression of which can also be monitored by fluorescence of the H2B-mIFP fusion. The size of each element is drawn approximately to scale.

**Figure 1 - figure supplement 2.**
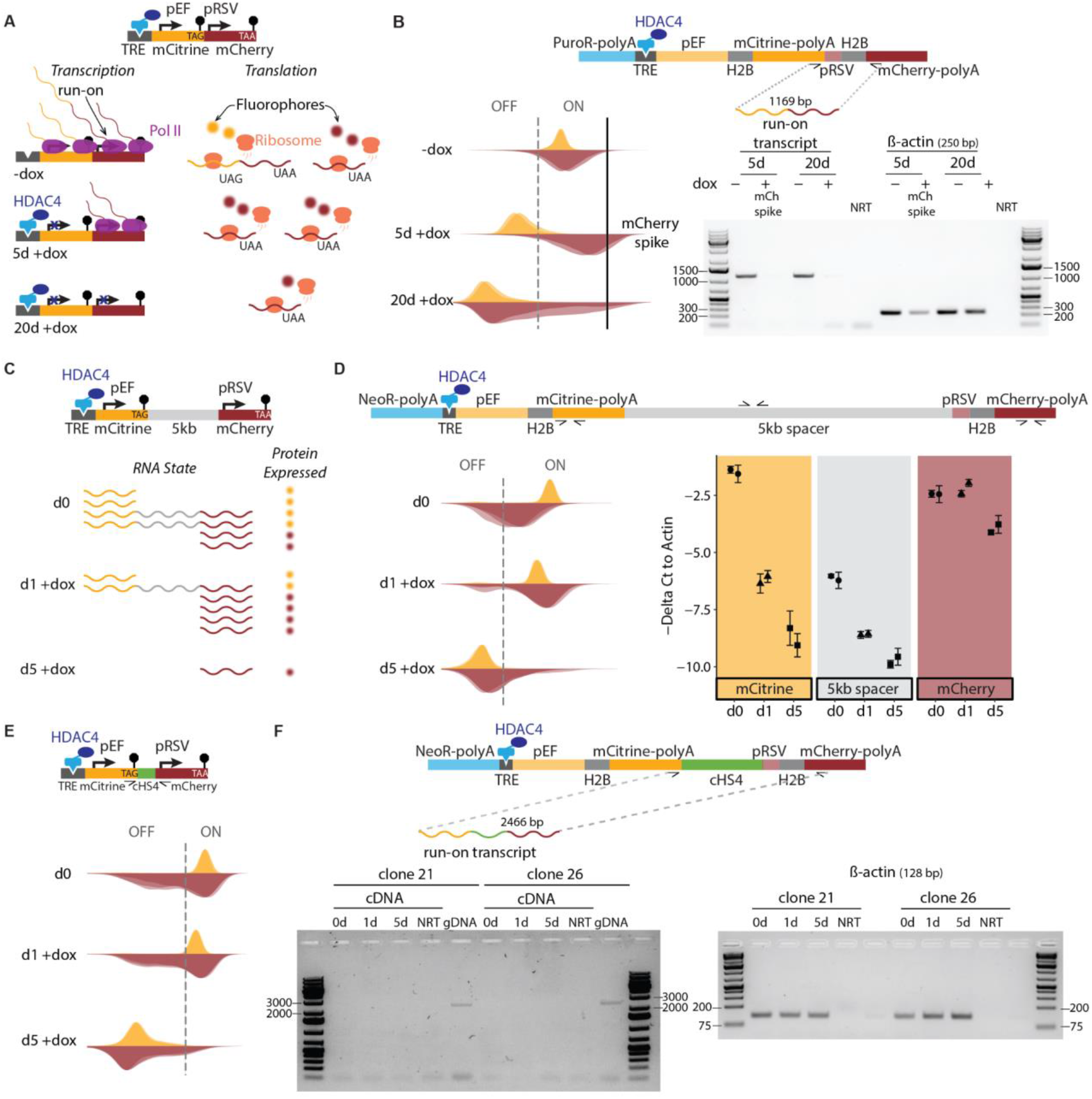
Transcriptional run-on from pEF over pRSV. **(A)** Schematic representation of transcription (left) and translation (right) of NS reporter (top) in K562 during HDAC4 recruitment. Transcriptional run-on from pEF into pRSV occurs despite the presence of polyA (-dox row, left). Note that mCitrine and mCherry genes both have stop codons at their 3’ end (black stop signs) and neither gene has an IRES at the 5’ end, such that mCherry can only be translated from transcripts initiated at pRSV (-dox row, right). When pEF is silenced by HDAC4 for 5 days (5d +dox row, left), run-on no longer occurs and transcription initiation at the pRSV increases, resulting in more mCherry protein (5d +dox row, right). After 20 days of HDAC4 recruitment (20d +dox row), both genes are silenced. **(B)** (top) Schematic of the NS spreading reporter (as described in Figure S1), showing primers for amplifying run-on transcript. (Left) Fluorescence distribution of mCitrine and mCherry across timepoints of HDAC4 recruitment at the NS reporter in K562, matching schematics in (A) for each row. Flow cytometry data from 3 replicates of recruitment are shown as overlaid semi-transparent distributions. Vertical dashed line shows the threshold for ON/OFF populations. After 5 days of recruitment (5d +dox) cells were sorted for mCherry expression above those in the -dox sample (mCherry spike, to the right of black line). (Right) Agarose gel showing PCR products that use as template cDNA reverse transcribed from mRNA extracted from either: cells with no recruitment (-dox), cells sorted on high mCherry (mCherry spike on the left), or cells with 20 days of recruitment (20d +dox). PCR primers amplify either a region spanning across the mCitrine and mCherry genes (labelled 1169 bp run-on transcript at the top) or beta-actin as a control for cDNA input. **(C)** Schematic representation of mRNA and fluorescent proteins levels of 5kb reporter in CHO-K1 during HDAC4 recruitment. Transcription termination fails in mCitrine polyA causing Pol II to run-on into 5kb lambda spacer and mCherry gene without dox (d0), reducing initiation of transcription of mCherry until pEF is silenced at day 1 of dox (d1), allowing pRSV to increase expression of mCherry before being silenced by the chromatin regulator at day 5 of dox (d5). **(D)** (Top) Schematic of the 5kb spreading reporter (as described in Figure S1), showing the positions of the primers used for qPCR. (Left) Fluorescence distributions of reporter genes across timepoints of HDAC4 recruitment to 5kb reporter in CHO-K1. Flow cytometry data from independent clonal cell lines are shown as overlaid semi-transparent distributions. Vertical dashed lines show the threshold for ON/OFF populations. (Right) qPCR was performed on cDNA reverse transcribed from mRNA on regions of mCitrine, 5kb lambda spacer, and mCherry after 0, 1, and 5 days of HDAC4 recruitment. Delta Cts with respect to Beta-actin are shown for two clones used as biological replicates. Error bars are one standard deviation of three technical replicates within each clone. Beta-actin was used as a control for normalization. **(E)** (Top) Schematic of HDAC4 recruitment at a construct where the reporter genes are separated by a full cHS4 insulator (SH). (Bottom) Fluorescence distributions of reporter genes across timepoints of HDAC4 recruitment to the SH construct in CHO-K1. Flow cytometry data from 3 replicates of recruitment are shown as overlaid semi-transparent distributions. Vertical dashed line shows the threshold for ON/OFF populations. **(F)** (Top) Schematic of the SH spreading reporter, showing PCR primers that detect the mRNA produced if run-on occurs. (Bottom, left) Agarose gel showing PCR products for a region spanning across the mCitrine and mCherry genes including the cHS4 insulator was performed on cDNA from CHO-K1 HDAC4 SH cells in two clones at 0, 1 and 5 days of HDAC4 recruitment. Genomic DNA (gDNA) was used as a control for primer validation. (Bottom, right) Beta-actin was amplified as a control for cDNA input. **Figure 1 - figure supplement 2 – source data.** Original gel images from RT-PCR.

**Figure 2 - figure supplement 1.**
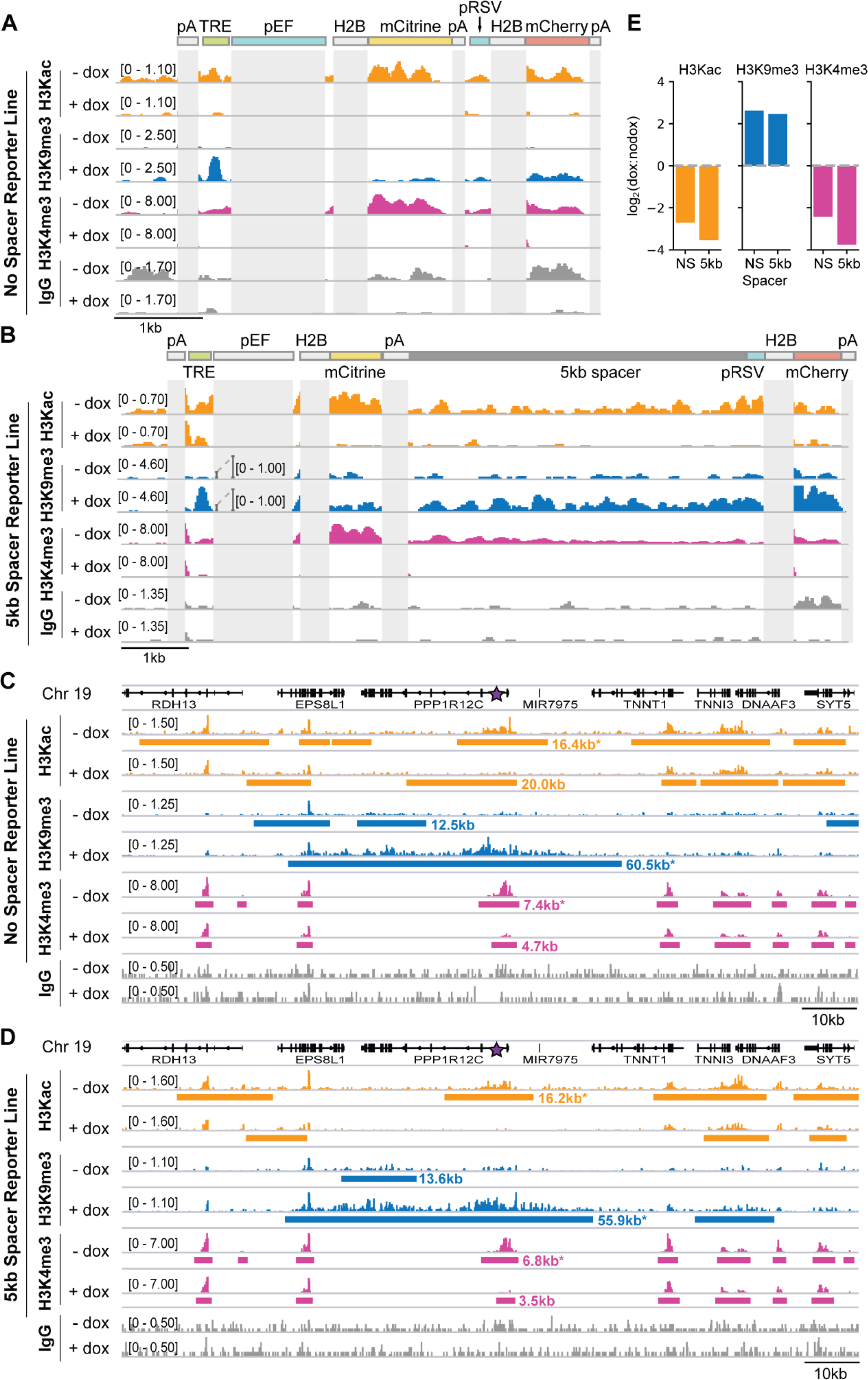
Changes in chromatin modifications at the two-gene reporter and surrounding AAVS1 locus after recruitment of KRAB for five days in K562 cells. **(A-B)** Genome browser tracks of histone 3 acetyl-lysine (H3Kac), histone 3 lysine 9 trimethylation (H3K9me3), histone 3 lysine 4 trimethylation (H3K4me3), and IgG, as measured by CUT&RUN in K562 cells with and without dox-mediated recruitment of rTetR-KRAB to the **(A)** no-spacer reporter and **(B)** 5kb reporter. Numbers in square brackets indicate y-axis (CPM) scaling. Reporter elements that also appear in the human genome or that are duplicated within the reporter (i.e. pEF, H2B, and polyA [pA]) are masked in light gray. **(C-D)** Genome browser tracks of H3Kac, H3K9me3, H3K4me3, and IgG ChromaBlocks-identified domains with and without recruitment of rTetR-KRAB, looking at the surrounding locus where the **(C)** NS reporter or **(D)** 5kb reporter is integrated in cells (purple star within first intron of PPP1R12C, which is oriented in the reverse direction). Note that this snapshot does not include an in situ representation of the reporter, which instead has been appended to the reference genome as a separate chromosome in order to preserve gene annotations. Domains for H3K9me3, H3Kac, and H3K4me3 are depicted as horizontal bars and were called with ChromaBlocks to estimate spreading distance with KRAB recruitment. Domains including or proximal to the integration site are annotated with their domain lengths (excludes the reporter length). **(E)** Quantification of modification level changes with KRAB recruitment. Coordinates of the ChromaBlocks-identified domains with asterisks in (C) and (D) were used to define a region for signal integration for the corresponding dox-treated (+dox) or untreated (-dox) sample. Log2 ratios of the total dox signal to total nodox signal within these regions are shown for each histone modification for each reporter cell line.

**Figure 2 - figure supplement 2.**
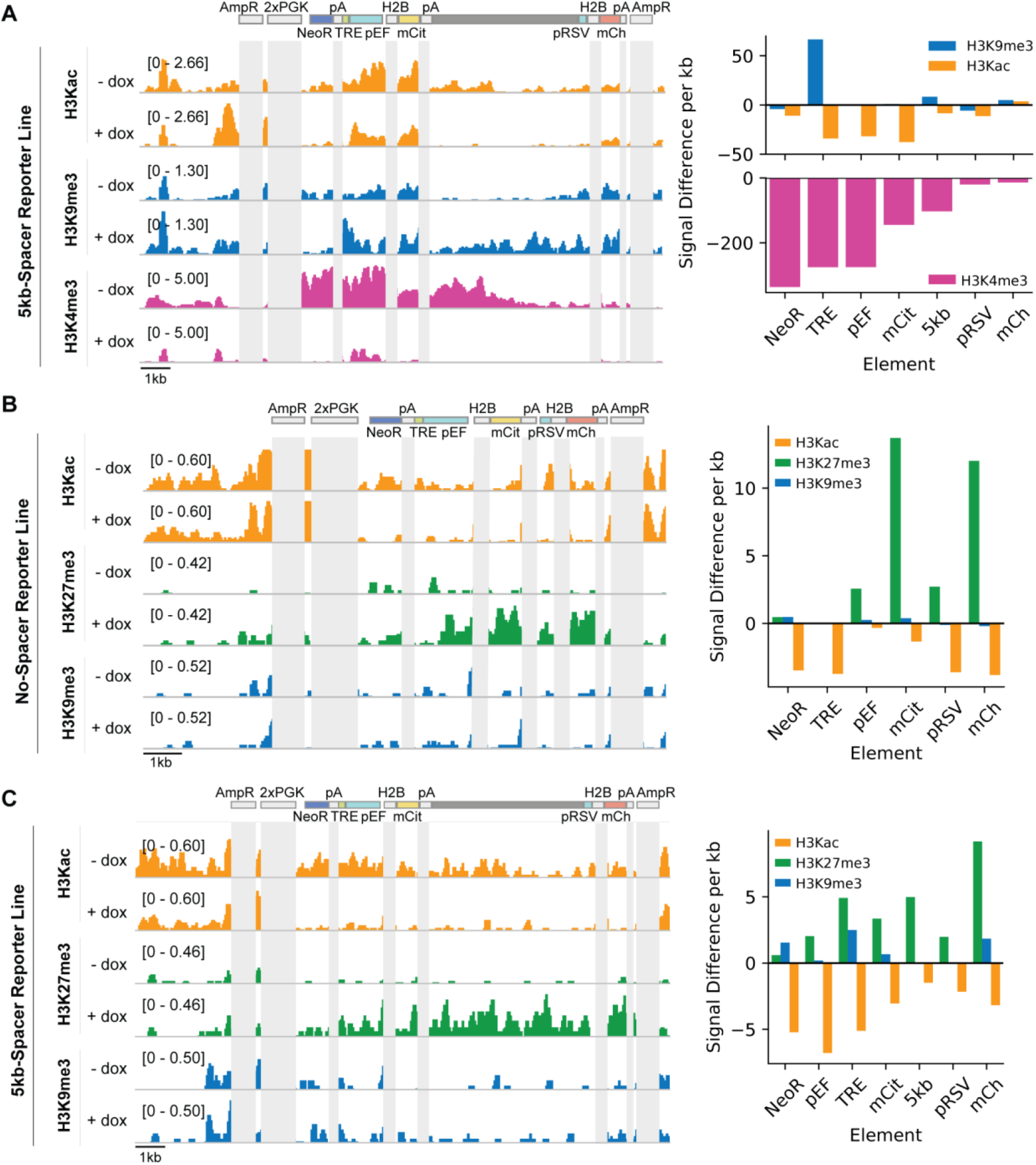
Recruitment of KRAB or HDAC4 for five days in CHO-K1 cells induces changes in chromatin modifications at the two-gene reporters. Genome browser tracks of histone 3 acetyl-lysine (H3Kac), histone 3 lysine 9 trimethylation (H3K9me3), histone 3 lysine 27 trimethylation (H3K27me3), and histone 3 lysine 4 trimethylation (H3K4me3) with (+dox) and without (-dox) recruitment of **(A)** rTetR-KRAB to the 5kb-spacer reporter, **(B)** rTetR-HDAC4 to the NS reporter and **(C)** rTetR-HDAC4 to the 5kb reporter. Reporter and MI-HAC elements that appear multiple times or share significant sequence identity (i.e. H2B, polyA [pA], PGK promoters, ampicillin resistance [AmpR]) are masked in light gray).

**Figure 3 - figure supplement 1.**
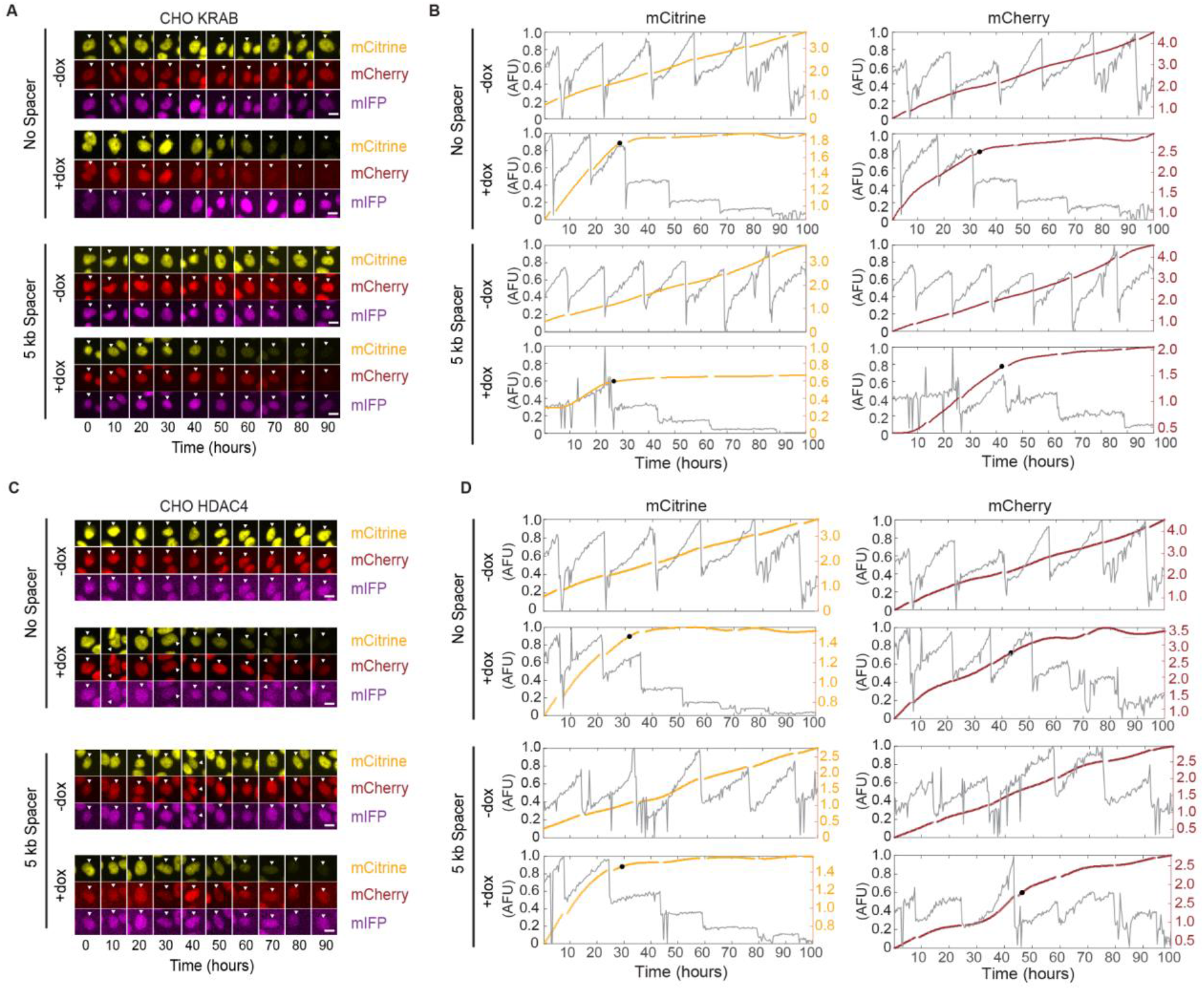
Example images and single-cell analysis of silencing dynamics from time-lapse microscopy of CHO-K1 cells. **(A, C)** Filmstrips at 10 hour intervals of representative tracked single-cells upon dox induced **(A)** KRAB and **(C)** HDAC4 silencing from time-lapse movies in all channels. Scale bar represents 10 microns. **(B, D)** Plot of nuclear intensity of the single-cell tracked in the filmstrip (A,C) in the mCitrine and mCherry channels. Grey traces represent the integrated nuclear intensity prior to stitching. The computationally stitched trace is shown in yellow (mCitrine), or red (mCherry) respectively. The black dot represents the time at which the cell was considered silenced.

**Figure 3 - figure supplement 2.**
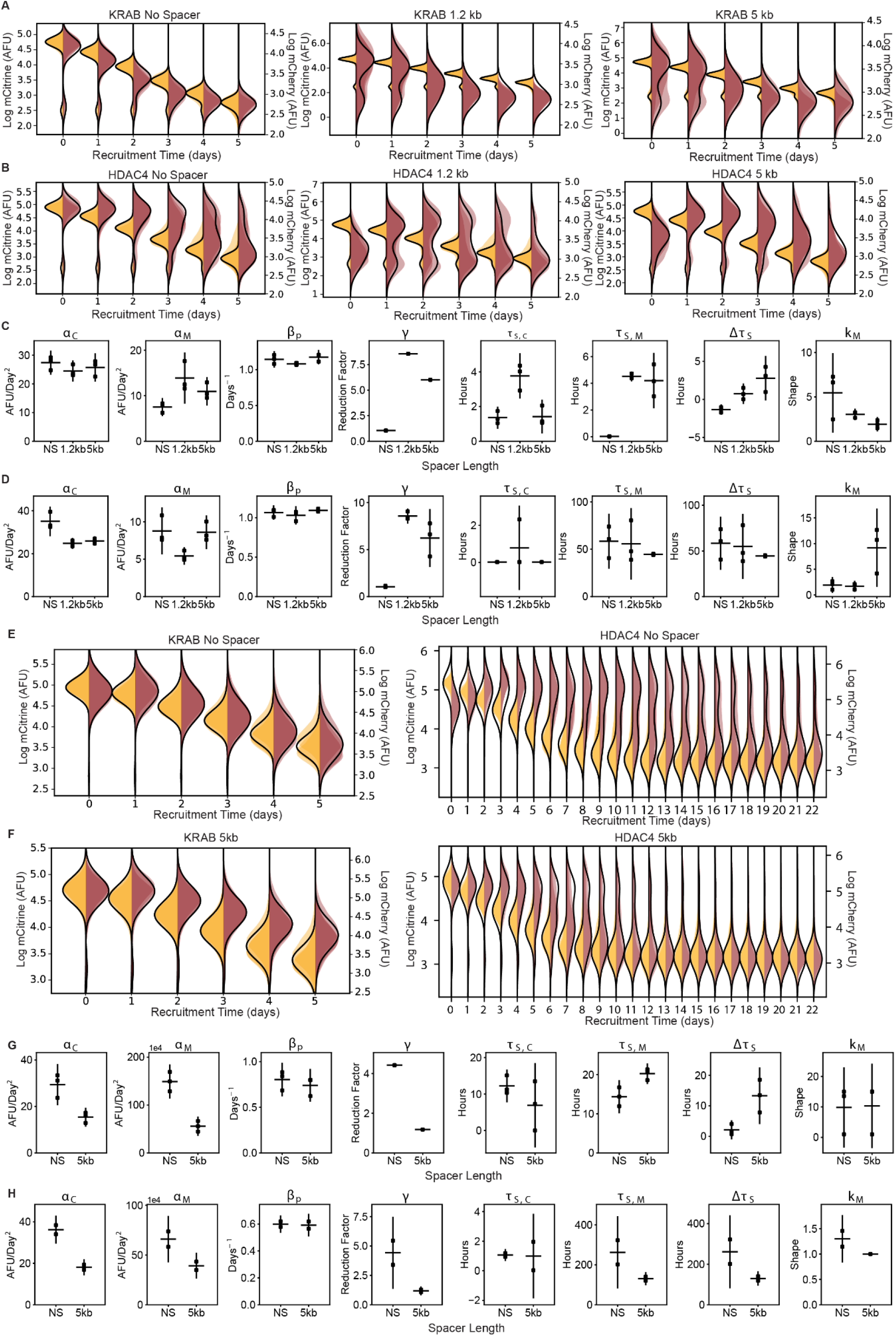
Dynamics of silencing measured by flow cytometry and fit by gene expression model. **(A-B)** Overlaid daily distributions of mCitrine (transparent yellow) and mCherry (transparent red) fluorescence from flow cytometry during recruitment of **(A)** KRAB and **(B)** HDAC4 in CHO-K1 for each spreading reporter indicated in the titles with average fit to the model in Fig 2F (black line) (n=3 clones). **(C-D)** Parameters from probabilistic model fit (Figure 3F) of flow cytometry data after **(C)** KRAB and **(D)** HDAC4 recruitment in CHO-K1. Each dot represents a clone of the given reporter, horizontal bar is mean delay, vertical bar is 90% confidence interval estimated using the t-distribution. **(E-F)** Overlaid replicates of daily distributions of mCitrine (transparent yellow) and mCherry (transparent red) fluorescence from flow cytometry, with average model fit (black line), during recruitment of KRAB (left, n=3) and HDAC4 (right, n=2) to **(E)** NS and **(F)** 5kb reporter in K562. **(G-H)** Parameters from probabilistic model fit of flow cytometry data after **(G)** KRAB and **(H)** HDAC4 recruitment in K562. Each dot represents a replicate of the given reporter line, horizontal bar is mean delay, vertical bar is 90% confidence interval estimated using the t-distribution.

**Figure 3 - figure supplement 3.**
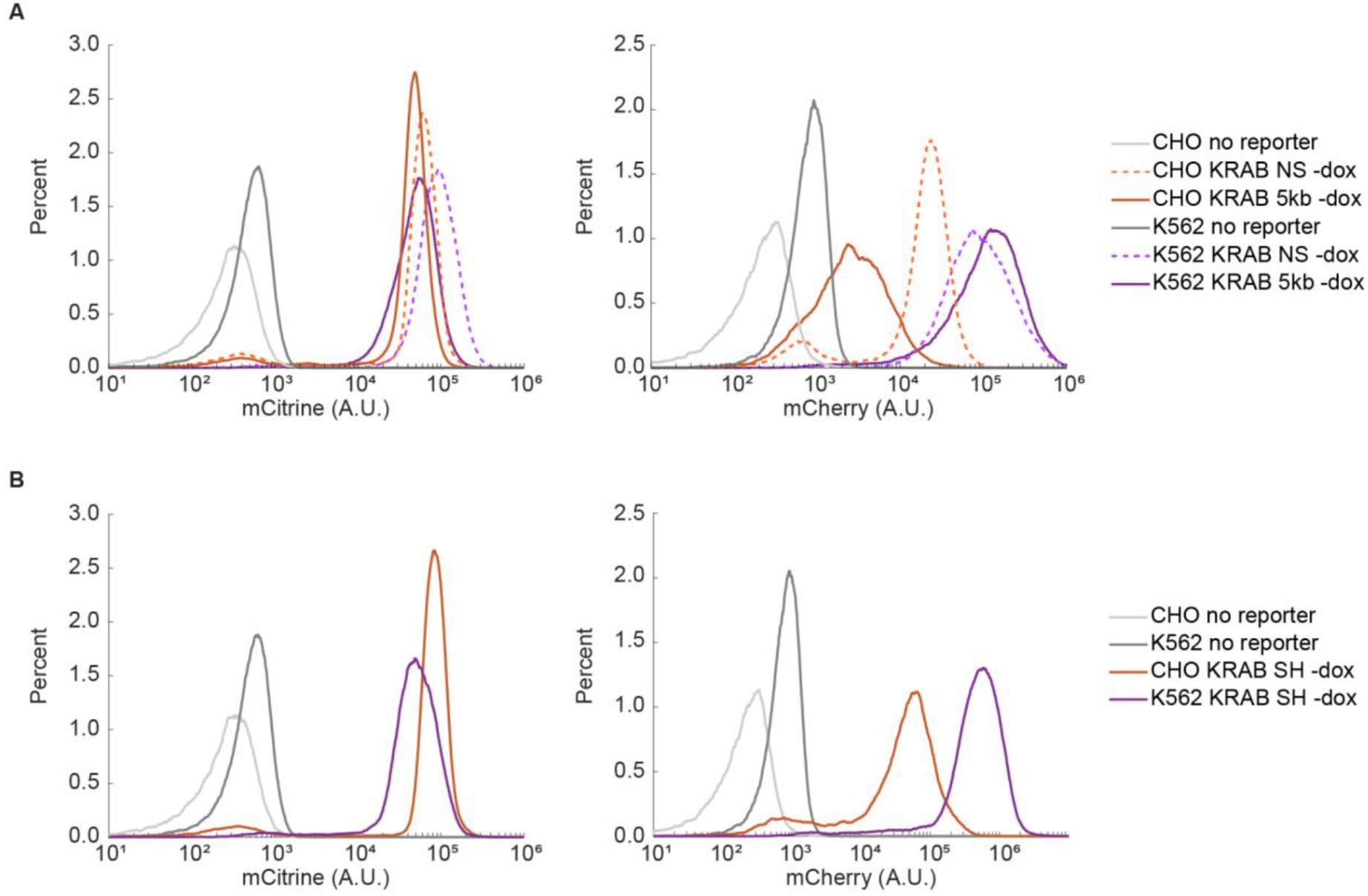
Steady-state expression in the absence of dox in CHO-K1 versus K562 cells. Fluorescence distributions of pEF-mCitrine (left) and pRSV-mCherry (right) from flow cytometry in the absence of dox-mediated recruitment in CHO-K1 (orange) versus K562 (purple) in different reporter constructs: **(A)** NS and 5kb, and **(B)** SH insulator. Note that pRSV-mCherry is always expressed at higher levels in K562 compared to CHO-K1.

**Figure 4 - figure supplement 1.**
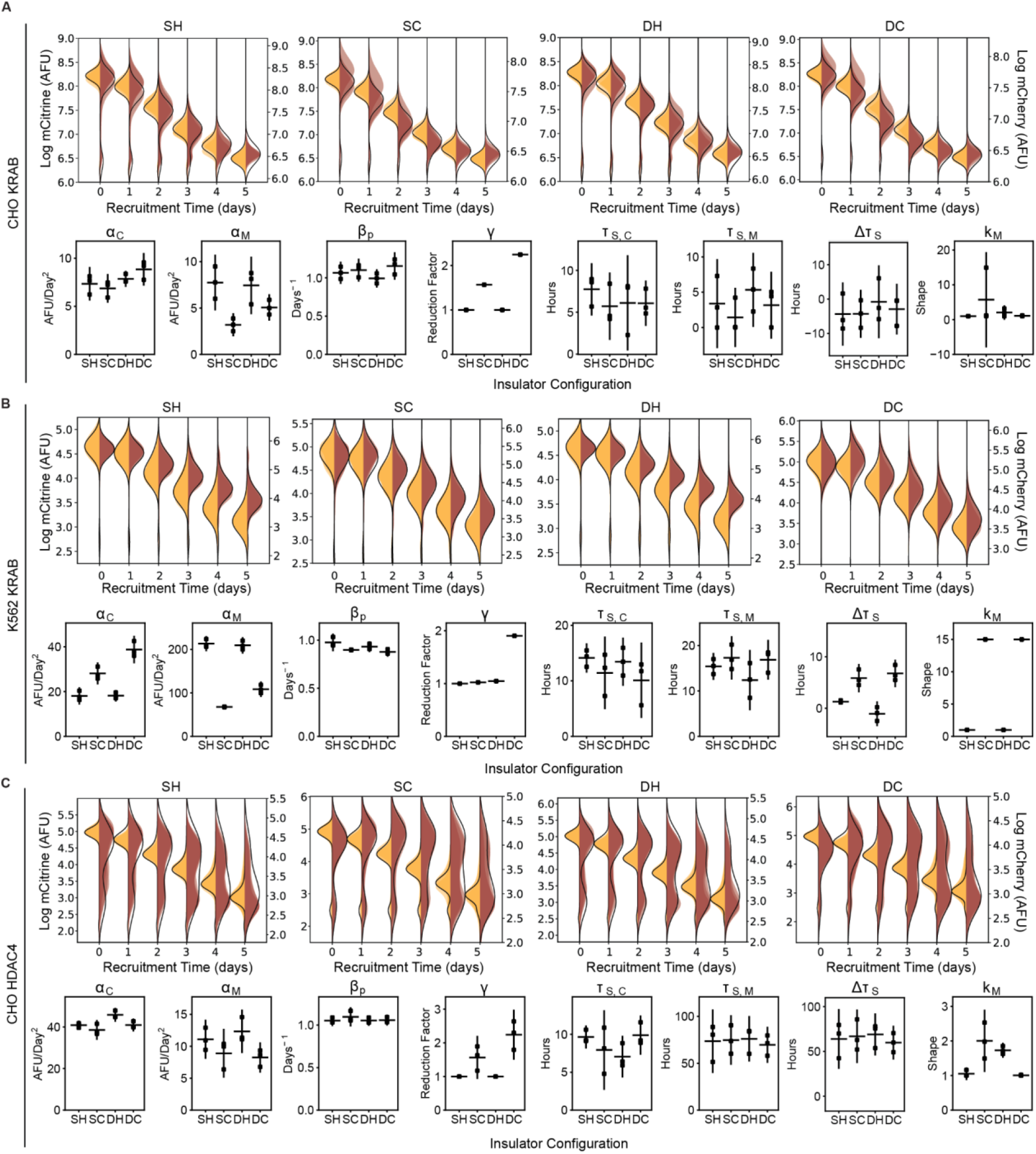
The effect of all insulator configurations on spreading dynamics of transcriptional silencing in CHO-K1 and K562. (Top panels of A-C) Overlaid replicates of daily distributions of mCitrine (transparent yellow) and mCherry (transparent red) fluorescence from flow cytometry and average model fits (black lines) during recruitment of : **(A)** KRAB in CHO-K1, **(B)** K562 with KRAB, and **(C)** CHO-K1 with HDAC4. (Bottom panels of A-C) Parameters from probabilistic model fit of flow cytometry data of insulator reporters with **(A)** KRAB recruitment in CHO-K1, **(B)** K562 with KRAB, and **(C)** CHO-K1 with HDAC4. Each dot represents a replicate of the given insulator reporter line, horizontal bar is mean delay, vertical bar is 90% confidence interval estimated using the t-distribution.

**Figure 4 - figure supplement 2.**
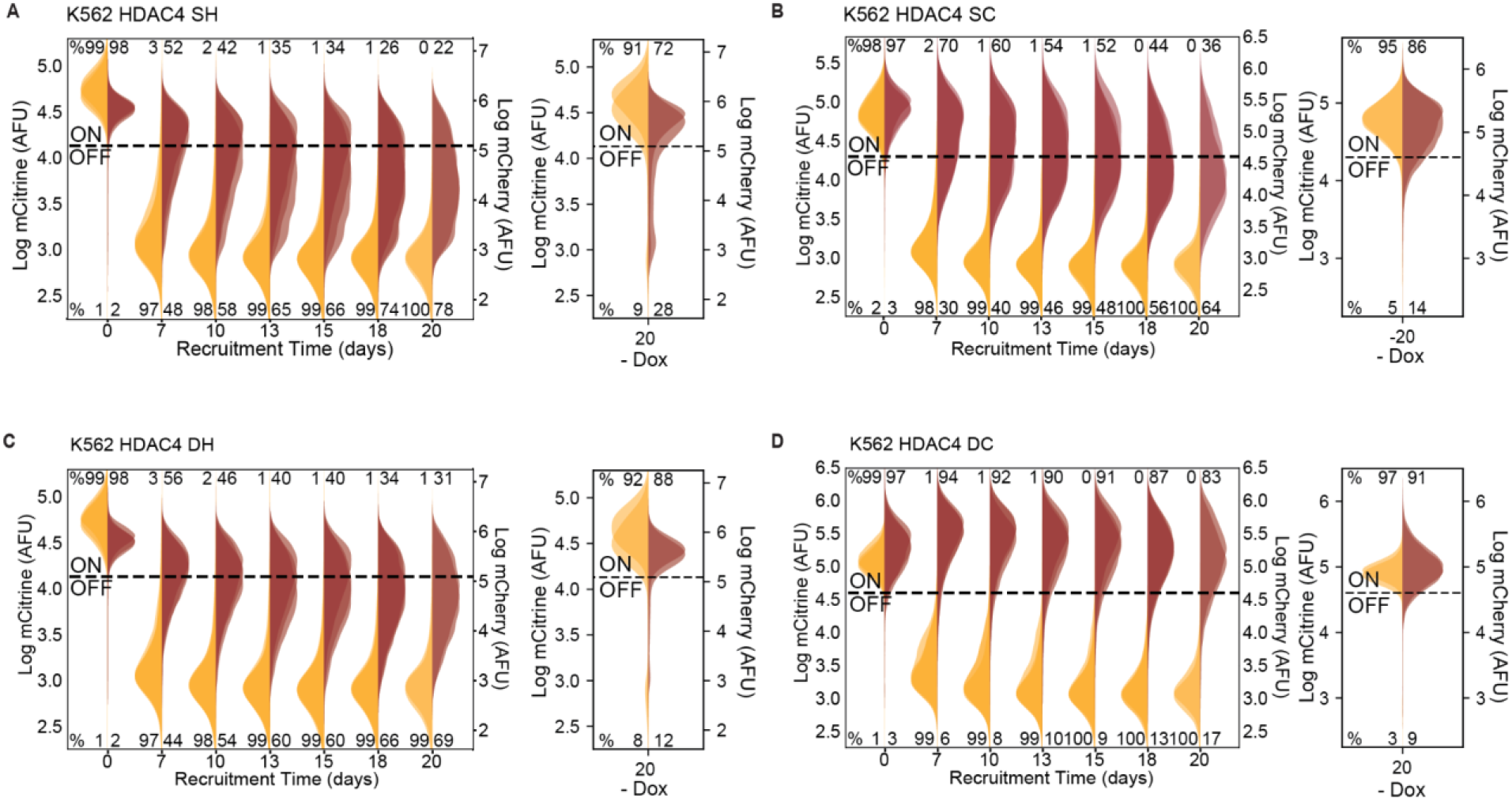
Insulators attenuate spreading of silencing via HDAC4 in K562. Overlaid replicates of distributions of mCitrine (transparent yellow) and mCherry (transparent red) fluorescence from flow cytometry data during extended recruitment of HDAC4 (left, n=3) and 20 days without recruitment (right, n=2) in K562 with insulator geometries defined in Figure 4A and B: **(A)** SH, **(B)** SC, **(C)** DC, and **(D)** DH.

**Figure 4 - figure supplement 3.**
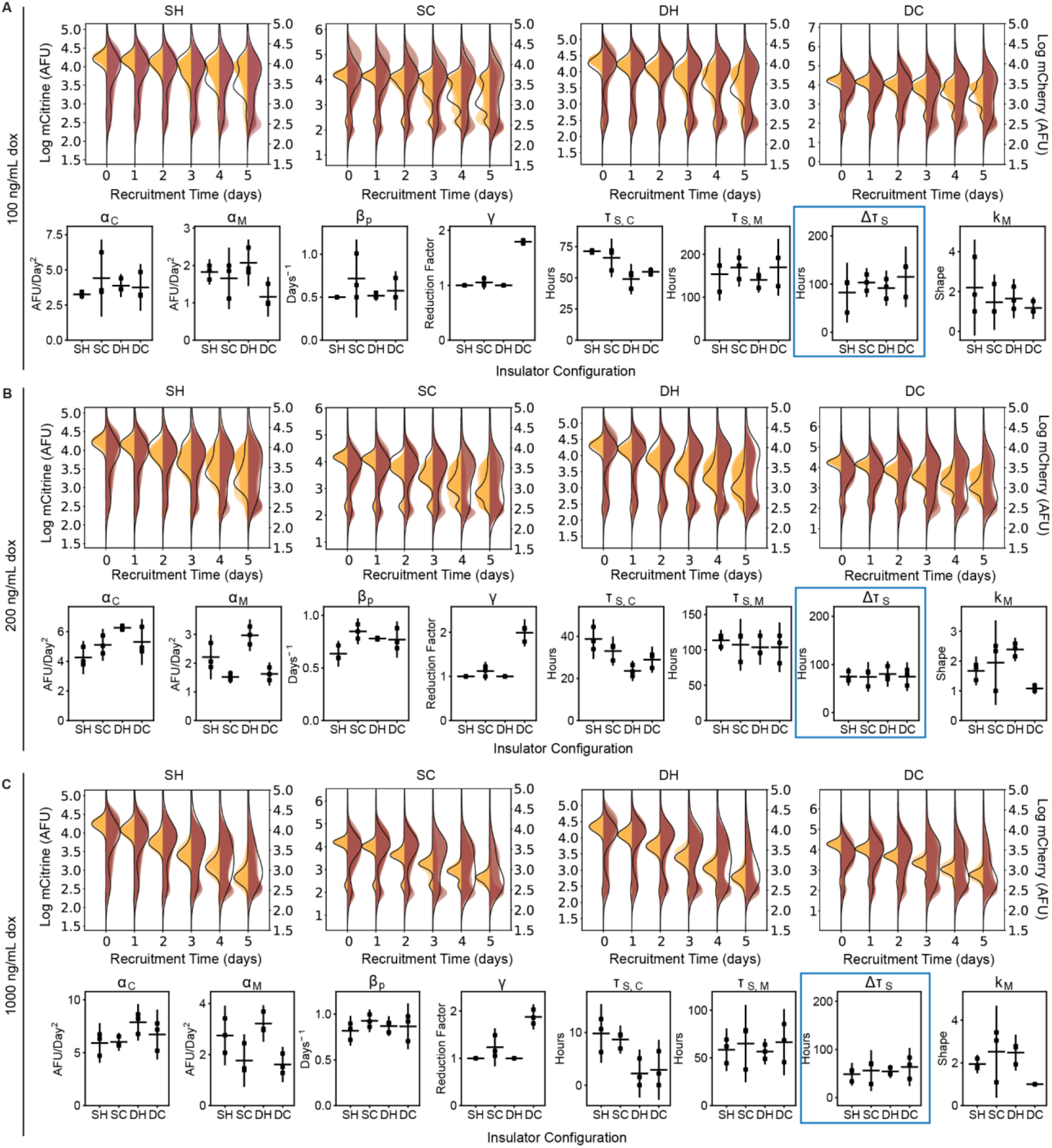
Dynamics of spreading upon weaker gene targeting insulator reporters with HDAC4 at lower dox concentrations in CHO-K1. (Top panels of A-C) Overlaid replicates of daily distributions of mCitrine (transparent yellow) and mCherry (transparent red) fluorescence from flow cytometry and average model fits (black lines) during recruitment of HDAC4 in CHO-K1 at **(A)** 100 ng/mL, **(B)** 200 ng/mL, and **(C)** 1000 ng/mL (saturating dox). (Bottom panels of A-C) Parameters from probabilistic model fit of flow cytometry data for insulator reporters with HDAC4 recruitment in CHO-K1 at: **(A)** 100 ng/mL, **(B)**200 ng/mL, and **(C)** and 1000 ng/mL. Each dot represents a replicate of the given insulator reporter line, horizontal bar is mean delay, vertical bar is 90% confidence interval estimated using the t-distribution (n=3). Silencing delay times (*Δτ*_*S*_) highlighted in blue.

**Figure 4 - figure supplement 4.**
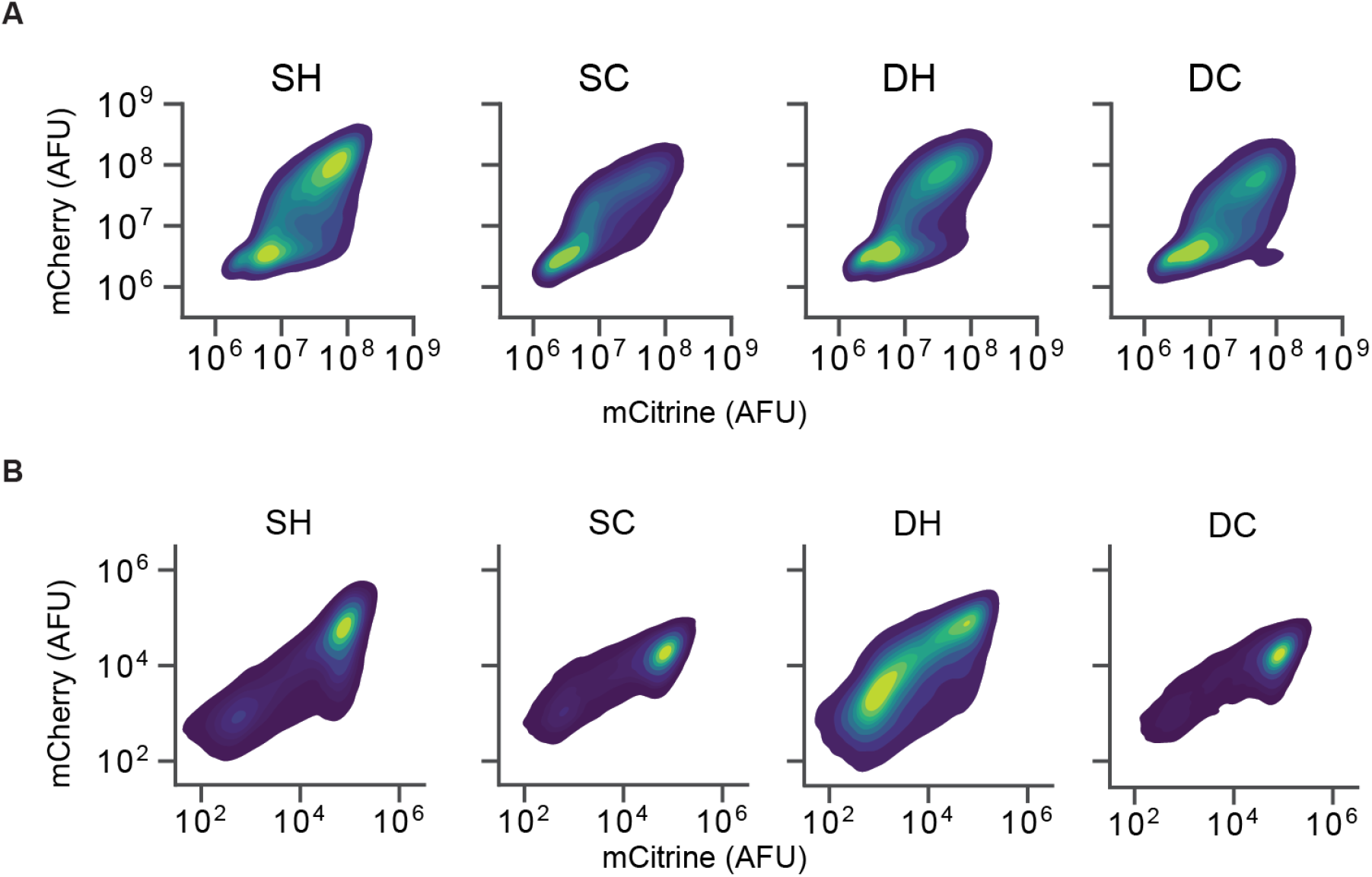
Insulators do not block spreading of silencing with weaker gene targeting at lower dox concentrations. 2D density plots of mCitrine and mCherry fluorescence measured by flow cytometry in CHO-K1 cells with insulator constructs indicated in the title of each graph after 5 days of **(A)** 200 ng/mL dox recruitment of HDAC4, or **(B)** 4 ng/mL dox recruitment of KRAB.

**Figure 4 - figure supplement 5.**
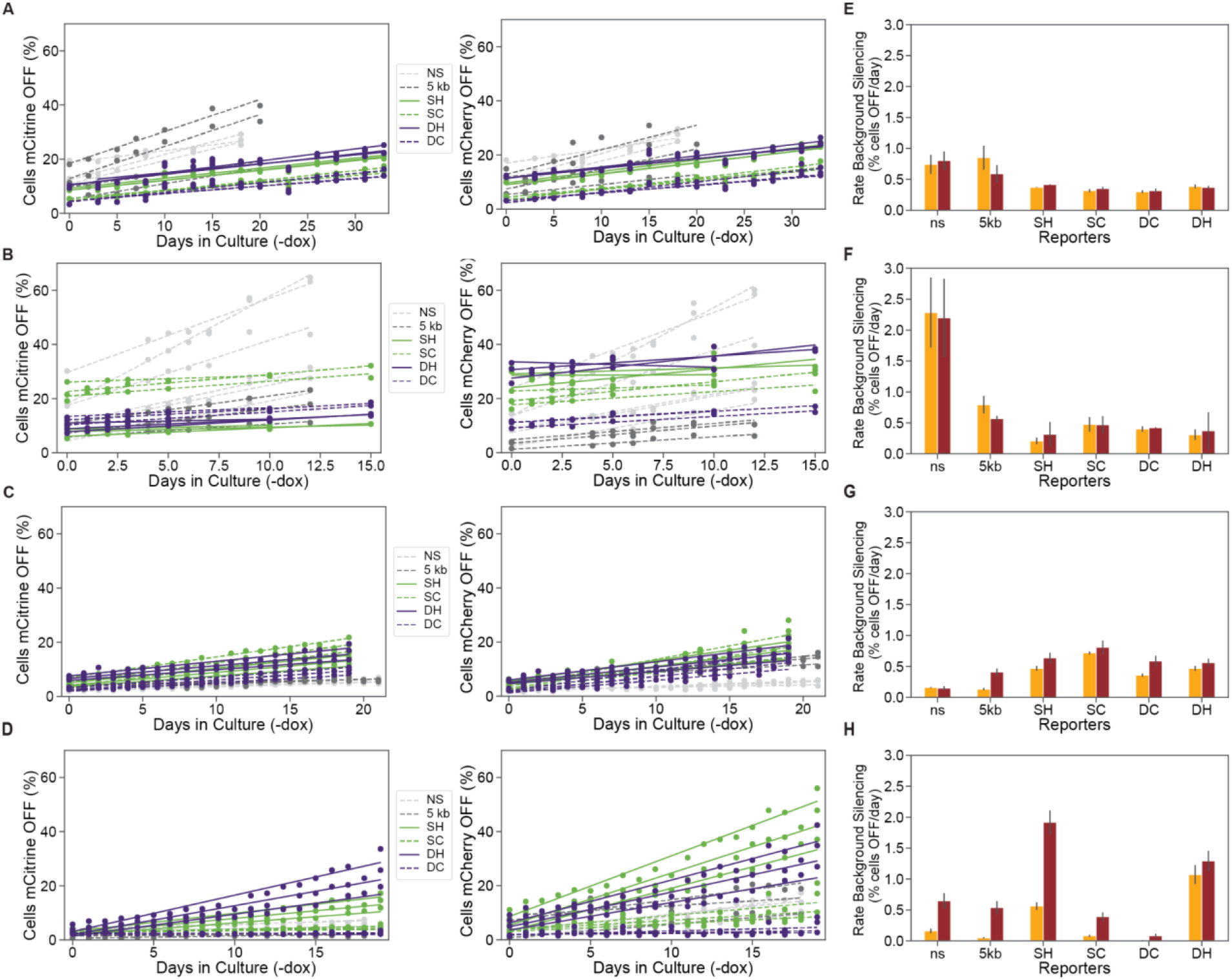
Insulators prevent background silencing of reporter genes. **(A-D)** Background levels of mCitrine (left) and mCherry silencing (right) in reporters with lambda spacers or insulators in the absence of dox for: **(A)** CHO-K1 KRAB, **(B)** CHO-K1 HDAC4, **(C)** K562 KRAB, and **(D)** K562 HDAC4 cell lines. **(E-H)** Rates of background silencing of mCitrine and mCherry in the absence of dox in: **(E)** CHO-K1 KRAB, **(F)** CHO-K1 HDAC4, **(G)** K562 KRAB, and **(H)** K562 HDAC4 cell lines. Error bars represent standard error of mean for 3 clones or replicates.

**Figure 5 - figure supplement 1.**
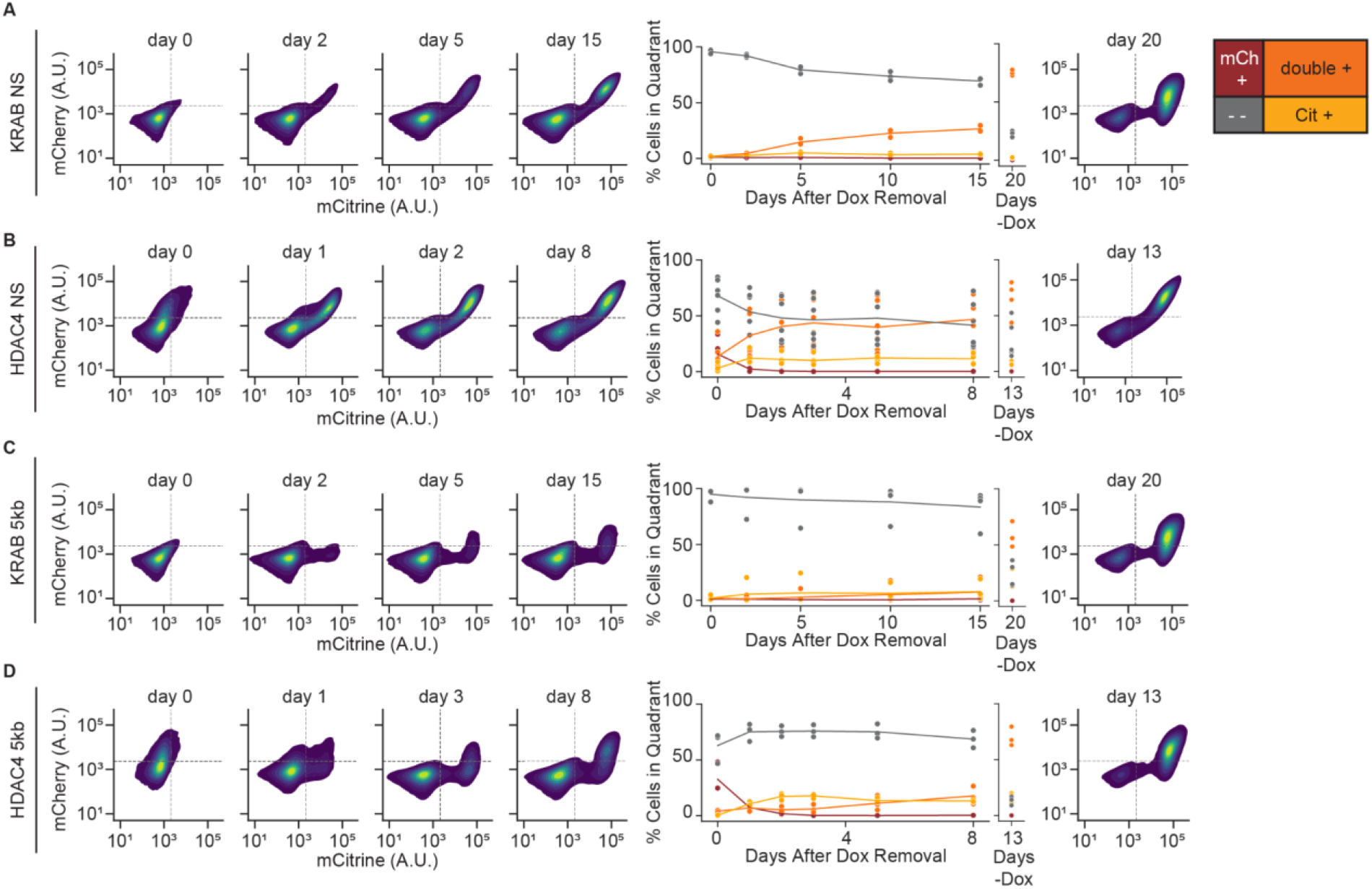
Reactivation of gene expression in CHO-K1 NS and 5kb reporter lines. (Left) 2D density plots of mCitrine and mCherry fluorescence from flow cytometry at different timepoints of reactivation (day 0 represents the end of 5 days of CR recruitment), (middle) percentages of cells in each quadrant as a function of release time, and (right) 2D density plots for no dox controls in: **(A)** NS reporter with KRAB (n=3 clones), **(B)** NS reporter with HDAC4 (n=6 clones), **(C)** 5kb reporter with KRAB (n=4 clones), and **(D)** 5kb reporter with HDAC4 (n=3 clones).

**Figure 5 - figure supplement 2.**
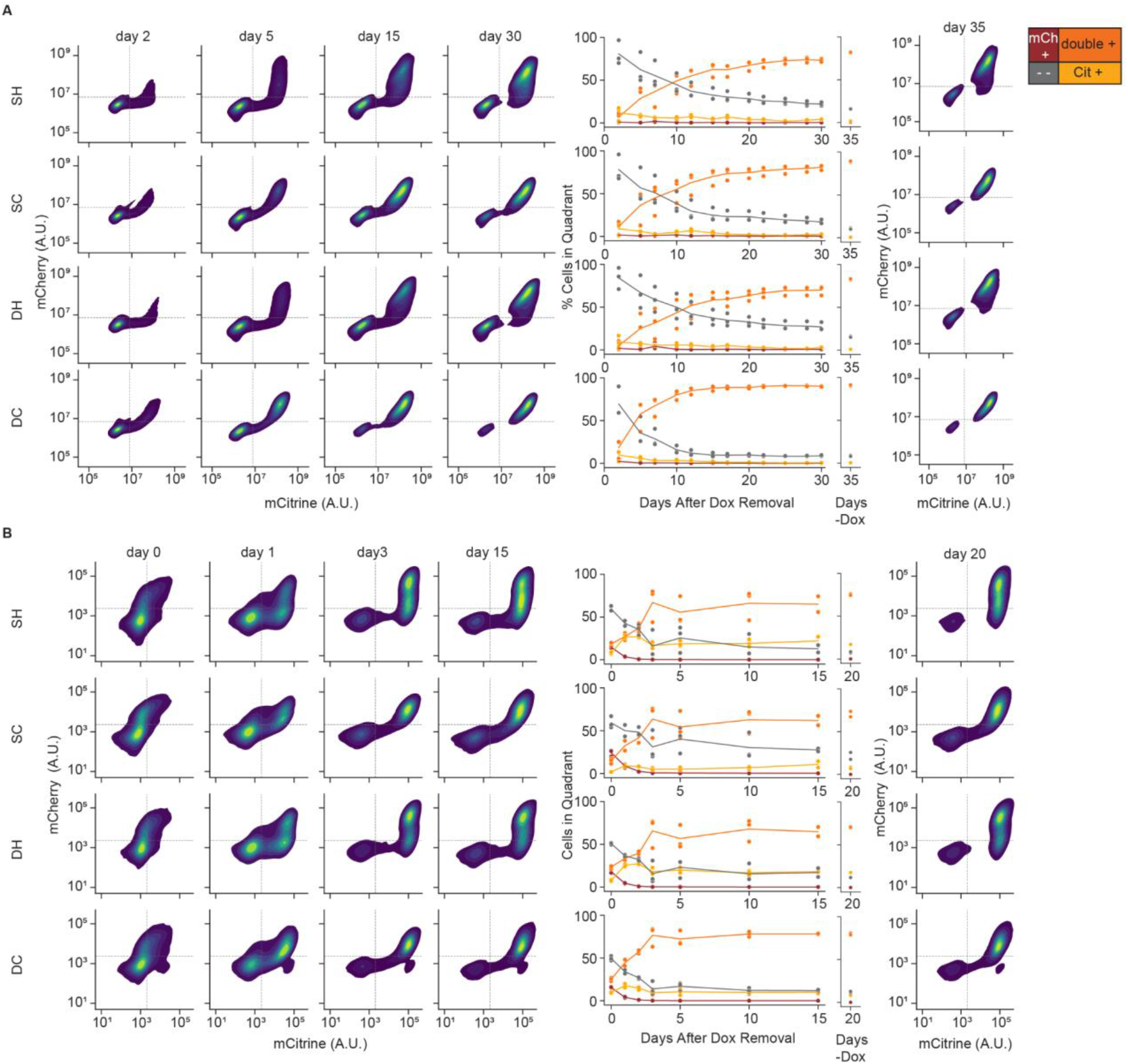
Reactivation of gene expression in CHO-K1 insulator reporter lines. (Left) 2D density plots of mCitrine and mCherry fluorescence from flow cytometry at different timepoints of reactivation (day 0 represents the end of 5 days of CR recruitment), (middle) percentages of cells in each quadrant as a function of release time, and (right) 2D density plots for no dox controls in: **(A)** insulators with KRAB and **(B)** insulators with HDAC4. Replicates are from biological replicates of multiclonal populations (n=3).

**Figure 5 - figure supplement 3.**
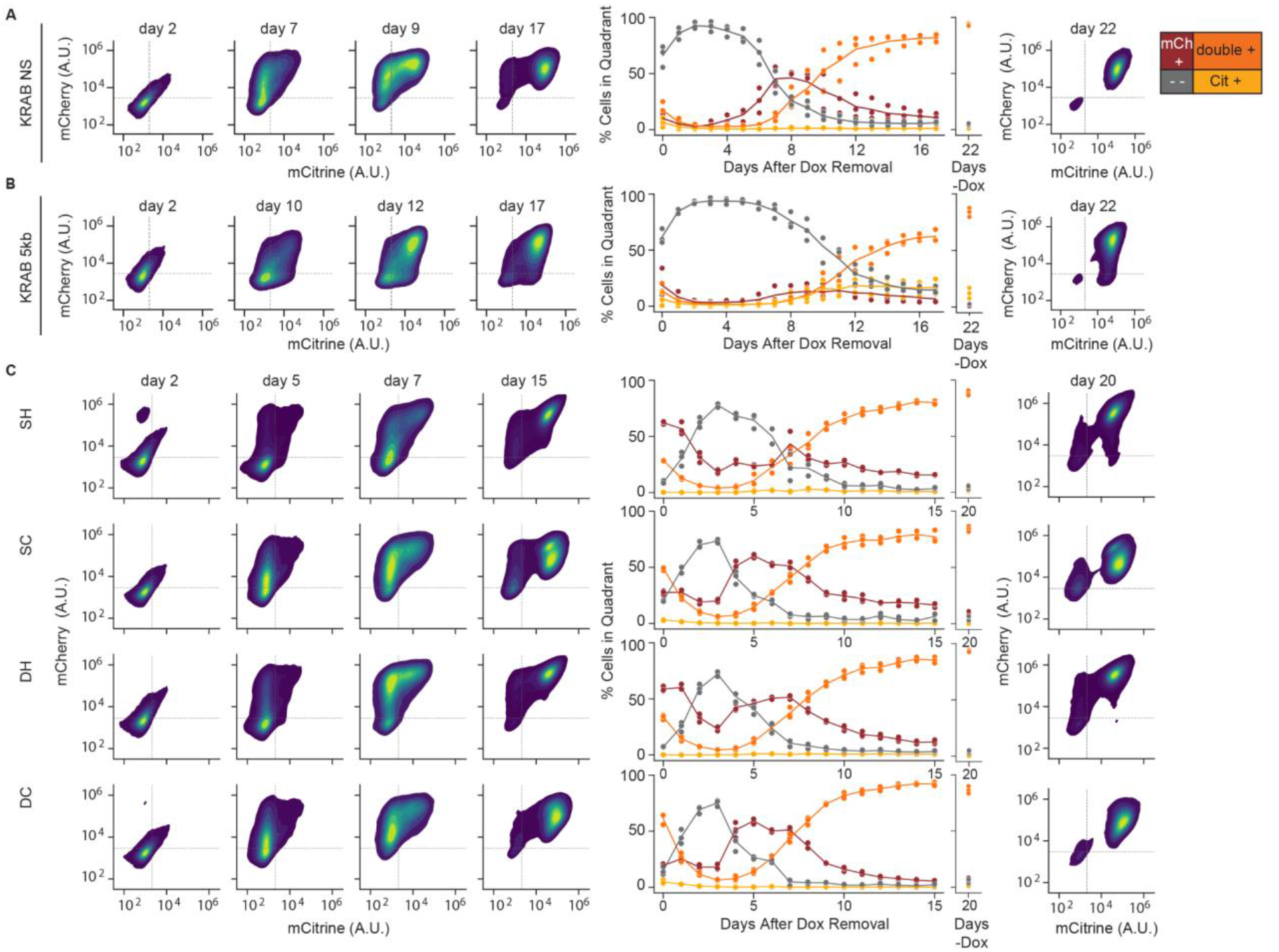
Reactivation of gene expression in K562 cell reporter lines. (Left) 2D density plots of mCitrine and mCherry fluorescence from flow cytometry at different timepoints of reactivation (day 0 represents the end of 5 days of CR recruitment), (middle) percentages of cells in each quadrant as a function of release time, and (right) 2D density plots for no dox controls in: **(A)** NS, **(B)** 5kb, and **(C)** insulator reporters after release of KRAB. Replicates are from biological replicates of multiclonal populations (n=3).

**Table S1.**
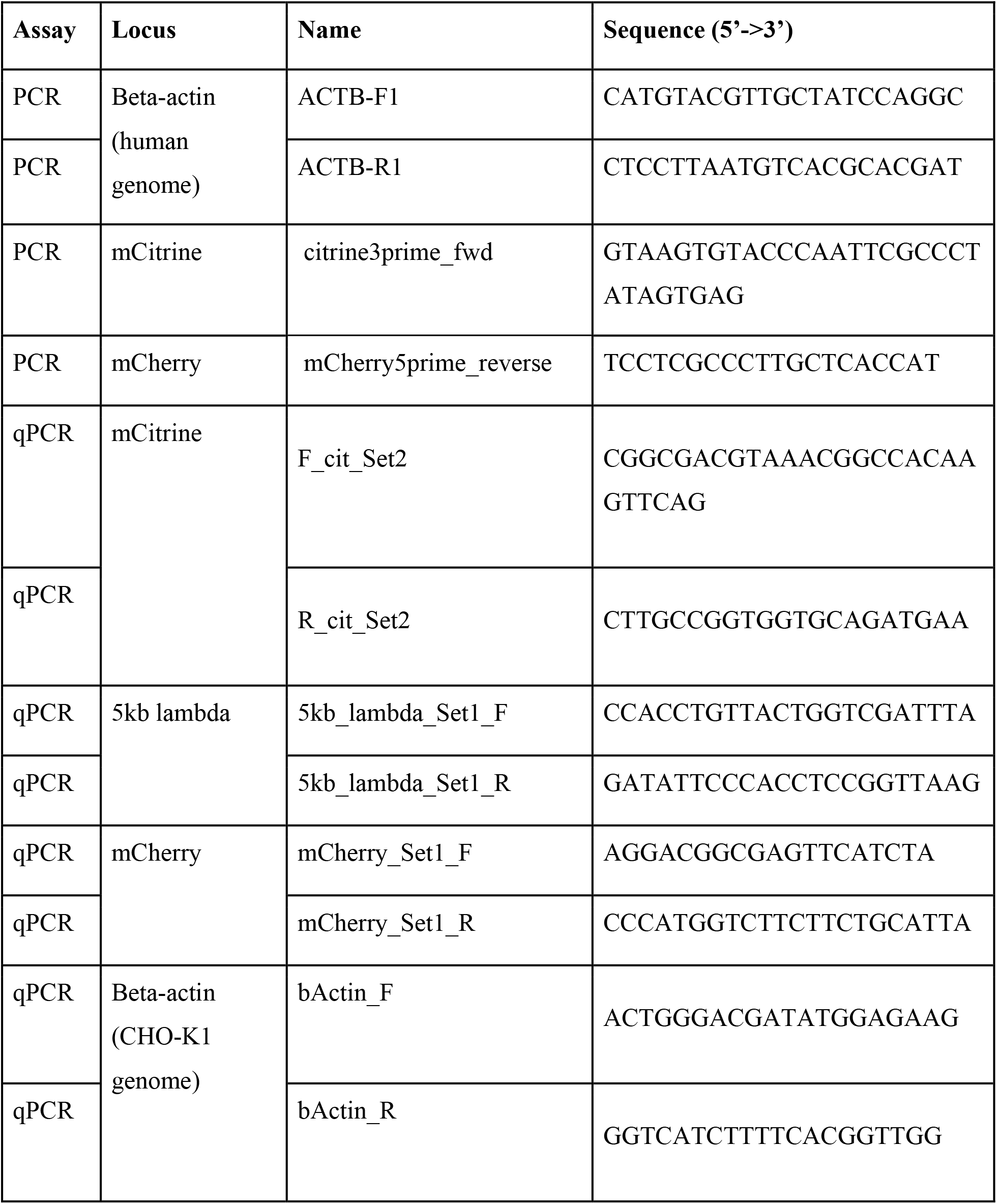
Primers used for investigating transcriptional interference.

**Table S2.** Delay times between mCitrine and mCherry silencing from model fit to flow cytometry data. The 90% confidence interval (CI) was estimated using the t-distribution.

| | | Reporter | $\Delta\tau_s$<br>(hour) | 90%<br>CI |
| --- | --- | --- | --- | --- |
| CHO-K1 | KRAB | NS | -1.34 | $\pm 0.59$ |
| | | 1.2 kb | 0.75 | $\pm 1.26$ |
| | | 5kb | 2.78 | $\pm 2.82$ |
| | | SH | -4.36 | $\pm 8.96$ |
| | | SC | -4.29 | $\pm 6.75$ |
| | | DH | -0.78 | $\pm 10.36$ |
| | | DC | -2.92 | $\pm 7.14$ |
| | HDAC4 | NS | 58.4 | $\pm 28.2$ |
| | | 1.2 kb | 54.9 | $\pm 34.8$ |
| | | 5kb | 44.5 | $\pm 1.1$ |
| | | SH | 64 | $\pm 32.5$ |
| | | SC | 67 | $\pm 28.9$ |
| | | DH | 69.1 | $\pm 22.6$ |
| | | DC | 59.9 | $\pm 17.9$ |
| K562 | KRAB | NS | 2.1 | $\pm 2.91$ |
| | | 5kb | 13.3 | $\pm 9.1$ |
| | | SH | 1.29 | $\pm 0.37$ |
| | | SC | 5.86 | $\pm 2.58$ |
| | | DH | -1.06 | $\pm 2.25$ |
| | | DC | 6.76 | $\pm 2.52$ |
| | HDAC4 | NS | 261 | $\pm 177$ |
| | | 5kb | 130 | $\pm 32$ |

**Movie S1. Spreading of silencing movie for KRAB NS.**

**Movie S2. Spreading of silencing movie for KRAB 5kb.**

**Movie S3. Spreading of silencing movie for HDAC4 NS.**

**Movie S4. Spreading of silencing movie for HDAC4 5kb.**

Movies S1-4 are zoomed-in views of spreading of silencing for KRAB and HDAC4 in the NS and 5kb reporters in CHO-K1. Each movie is digitally cropped to follow a subset of cells captured in the full frame. mCitrine fluorescence is pseudo-colored as yellow, and mCherry fluorescence is pseudo-colored in red. Timestamps are in HH:MM relative to dox addition at 0 hours.

## References

1. Bannister, A. J. & Kouzarides, T. Regulation of chromatin by histone modifications. Cell Res. 21, 381–395 (2011).

2. Zhang, P., Torres, K., Liu, X., Liu, C.-G. & Pollock, R. E. An Overview of Chromatin-Regulating Proteins in Cells. Curr. Protein Pept. Sci. 17, 401–410 (2016).

3. Jambhekar, A., Dhall, A. & Shi, Y. Roles and regulation of histone methylation in animal development. Nat. Rev. Mol. Cell Biol. 20, 625–641 (2019).

4. Völker-Albert, M., Bronkhorst, A., Holdenrieder, S. & Imhof, A. Histone Modifications in Stem Cell Development and Their Clinical Implications. Stem Cell Reports 15, 1196–1205 (2020).

5. Zhao, Z. & Shilatifard, A. Epigenetic modifications of histones in cancer. Genome Biol. 20, 245 (2019).

6. Keung, A. J., Joung, J. K., Khalil, A. S. & Collins, J. J. Chromatin regulation at the frontier of synthetic biology. Nat. Rev. Genet. 16, 159–171 (2015).

7. Thakore, P. I., Black, J. B., Hilton, I. B. & Gersbach, C. A. Editing the epigenome: technologies for programmable transcription and epigenetic modulation. Nat. Methods 13, 127–137 (2016).

8. Bintu, L. et al. Dynamics of epigenetic regulation at the single-cell level. Science 351, 720–724 (2016).

9. Ayyanathan, K. et al. Regulated recruitment of HP1 to a euchromatic gene induces mitotically heritable, epigenetic gene silencing: a mammalian cell culture model of gene variegation. Genes Dev. 17, 1855–1869 (2003).

10. Uckelmann, M. & Davidovich, C. Not just a writer: PRC2 as a chromatin reader. Biochem. Soc. Trans. 49, 1159–1170 (2021).

11. Zhang, T., Cooper, S. & Brockdorff, N. The interplay of histone modifications – writers that read. EMBO Rep. 16, 1467–1481 (2015).

12. Chung, J. H., Whiteley, M. & Felsenfeld, G. A 5’ element of the chicken beta-globin domain serves as an insulator in human erythroid cells and protects against position effect in Drosophila. Cell 74, 505–514 (1993).

13. Chung, J. H., Bell, A. C. & Felsenfeld, G. Characterization of the chicken beta-globin insulator. Proc. Natl. Acad. Sci. U. S. A. 94, 575–580 (1997).

14. Guye, P., Li, Y., Wroblewska, L., Duportet, X. & Weiss, R. Rapid, modular and reliable construction of complex mammalian gene circuits. Nucleic Acids Res. 41, e156 (2013).

15. Margolin, J. F. et al. Krüppel-associated boxes are potent transcriptional repression domains. Proc. Natl. Acad. Sci. U. S. A. 91, 4509–4513 (1994).

16. Deuschle, U., Meyer, W. K. & Thiesen, H. J. Tetracycline-reversible silencing of eukaryotic promoters. Mol. Cell. Biol. 15, 1907–1914 (1995).

17. Cong, L., Zhou, R., Kuo, Y.-C., Cunniff, M. & Zhang, F. Comprehensive interrogation of natural TALE DNA-binding modules and transcriptional repressor domains. Nat. Commun. 3, 968 (2012).

18. Gilbert, L. A. et al. CRISPR-mediated modular RNA-guided regulation of transcription in eukaryotes. Cell 154, 442–451 (2013).

19. Groner, A. C. et al. KRAB-zinc finger proteins and KAP1 can mediate long-range transcriptional repression through heterochromatin spreading. PLoS Genet. 6, e1000869 (2010).

20. Amabile, A. et al. Inheritable Silencing of Endogenous Genes by Hit-and-Run Targeted Epigenetic Editing. Cell 167, 219–232.e14 (2016).

21. O’Geen, H. et al. Ezh2-dCas9 and KRAB-dCas9 enable engineering of epigenetic memory in a context-dependent manner. Epigenetics Chromatin 12, 26 (2019).

22. Gjaltema, R. A. F. et al. KRAB-Induced Heterochromatin Effectively Silences PLOD2 Gene Expression in Somatic Cells and is Resilient to TGFβ1 Activation. Int. J. Mol. Sci. 21, (2020).

23. Groner, A. C. et al. The Krüppel-associated box repressor domain can induce reversible heterochromatization of a mouse locus in vivo. J. Biol. Chem. 287, 25361–25369 (2012).

24. Miska, E. A. et al. HDAC4 deacetylase associates with and represses the MEF2 transcription factor. EMBO J. 18, 5099–5107 (1999).

25. Wang, A. H. et al. HDAC4, a human histone deacetylase related to yeast HDA1, is a transcriptional corepressor. Mol. Cell. Biol. 19, 7816–7827 (1999).

26. Belozerov, V. E., Majumder, P., Shen, P. & Cai, H. N. A novel boundary element may facilitate independent gene regulation in the Antennapedia complex of Drosophila. EMBO J. 22, 3113–3121 (2003).

27. Yamaguchi, S. et al. A method for producing transgenic cells using a multi-integrase system on a human artificial chromosome vector. PLoS One 6, e17267 (2011).

28. Hockemeyer, D. et al. Genetic engineering of human pluripotent cells using TALE nucleases. Nat. Biotechnol. 29, 731–734 (2011).

29. Shearwin, K. E., Callen, B. P. & Egan, J. B. Transcriptional interference--a crash course. Trends Genet. 21, 339–345 (2005).

30. Skene, P. J., Henikoff, J. G. & Henikoff, S. Targeted in situ genome-wide profiling with high efficiency for low cell numbers. Nat. Protoc. 13, 1006–1019 (2018).

31. Meers, M. P., Bryson, T. D., Henikoff, J. G. & Henikoff, S. Improved CUT&RUN chromatin profiling tools. Elife 8, (2019).

32. Barkess, G. & West, A. G. Chromatin insulator elements: establishing barriers to set heterochromatin boundaries. Epigenomics 4, 67–80 (2012).

33. West, A. G., Huang, S., Gaszner, M., Litt, M. D. & Felsenfeld, G. Recruitment of histone modifications by USF proteins at a vertebrate barrier element. Mol. Cell 16, 453–463 (2004).

34. Recillas-Targa, F. et al. Position-effect protection and enhancer blocking by the chicken beta-globin insulator are separable activities. Proc. Natl. Acad. Sci. U. S. A. 99, 6883–6888 (2002).

35. Gowher, H., Brick, K., Camerini-Otero, R. D. & Felsenfeld, G. Vezf1 protein binding sites genome-wide are associated with pausing of elongating RNA polymerase II. Proc. Natl. Acad. Sci. U. S. A. 109, 2370–2375 (2012).

36. Yusufzai, T. M. & Felsenfeld, G. The 5’-HS4 chicken beta-globin insulator is a CTCF-dependent nuclear matrix-associated element. Proc. Natl. Acad. Sci. U. S. A. 101, 8620–8624 (2004).

37. Mutskov, V. J., Farrell, C. M., Wade, P. A., Wolffe, A. P. & Felsenfeld, G. The barrier function of an insulator couples high histone acetylation levels with specific protection of promoter DNA from methylation. Genes Dev. 16, 1540–1554 (2002).

38. Zhao, H. & Dean, A. An insulator blocks spreading of histone acetylation and interferes with RNA polymerase II transfer between an enhancer and gene. Nucleic Acids Res. 32, 4903–4919 (2004).

39. Pikaart, M. J., Recillas-Targa, F. & Felsenfeld, G. Loss of transcriptional activity of a transgene is accompanied by DNA methylation and histone deacetylation and is prevented by insulators. Genes Dev. 12, 2852–2862 (1998).

40. Tian, J. & Andreadis, S. T. Independent and high-level dual-gene expression in adult stem-progenitor cells from a single lentiviral vector. Gene Ther. 16, 874–884 (2009).

41. Thakore, P. I. et al. Highly specific epigenome editing by {CRISPR-Cas9} repressors for silencing of distal regulatory elements. Nat. Methods 12, 1143–1149 (2015).

42. Hathaway, N. A. et al. Dynamics and memory of heterochromatin in living cells. Cell 149, 1447–1460 (2012).

43. Dodd, I. B., Micheelsen, M. A., Sneppen, K. & Thon, G. Theoretical Analysis of Epigenetic Cell Memory by Nucleosome Modification. Cell 129, 813–822 (2007).

44. Hodges, C. & Crabtree, G. R. Dynamics of inherently bounded histone modification domains. Proceedings of the National Academy of Sciences 109, 13296–13301 (2012).

45. Erdel, F. & Greene, E. C. Generalized nucleation and looping model for epigenetic memory of histone modifications. Proc. Natl. Acad. Sci. U. S. A. 113, E4180–9 (2016).

46. Erdel, F. How Communication Between Nucleosomes Enables Spreading and Epigenetic Memory of Histone Modifications. Bioessays 39, 1–12 (2017).

47. Park, M., Patel, N., Keung, A. J. & Khalil, A. S. Engineering Epigenetic Regulation Using Synthetic Read-Write Modules. Cell vol. 176 227–238.e20 (2019).

48. Rudina, S. S. & Smolke, C. D. A Novel Chromatin-Opening Element for Stable Long-term Transgene Expression. bioRxiv 626713 (2019) doi:10.1101/626713.

49. Sprinzak, D. et al. Cis-interactions between Notch and Delta generate mutually exclusive signalling states. Nature 465, 86–90 (2010).

50. Sanjana, N. E. et al. A transcription activator-like effector toolbox for genome engineering. Nat. Protoc. 7, 171–192 (2012).

51. Oceguera-Yanez, F. et al. Engineering the AAVS1 locus for consistent and scalable transgene expression in human iPSCs and their differentiated derivatives. Methods 101, 43–55 (2016).

52. Janssens, D. & Henikoff, S. CUT&RUN: Targeted in situ genome-wide profiling with high efficiency for low cell numbers v3 (protocols.io.zcpf2vn). protocols.io (2019) doi:10.17504/protocols.io.zcpf2vn.

53. Hawkins, R. D. et al. Distinct epigenomic landscapes of pluripotent and lineage-committed human cells. Cell Stem Cell 6, 479–491 (2010).

54. Adhya, S. & Gottesman, M. Promoter occlusion: transcription through a promoter may inhibit its activity. Cell 29, 939–944 (1982).

55. Eszterhas, S. K., Bouhassira, E. E., Martin, D. I. K. & Fiering, S. Transcriptional interference by independently regulated genes occurs in any relative arrangement of the genes and is influenced by chromosomal integration position. Mol. Cell. Biol. 22, 469–479 (2002).

56. Yamaguchi, S. et al. A method for producing transgenic cells using a multi-integrase system on a human artificial chromosome vector. PLoS One 6, e17267 (2011).

